# Transcriptome analyses using regulon, functional category, and operon information with GINtool

**DOI:** 10.1101/2022.10.24.513545

**Authors:** Biwen Wang, Frans van der Kloet, Mariah B.M.J. Kes, Joen Luirink, Leendert W. Hamoen

## Abstract

When analysing transcriptome data, threshold values are chosen to decide whether the regulation of a gene is relevant or not, however this may result in the loss of valuable information. To overcome this problem it can be useful to analyse regulons instead of individual genes, to harness the statistical power of combining genes. Another advantage of a regulon-based analysis is that it provides direct insights into the activity of regulatory pathways, which is the essence of transcriptome analyses. We have developed a software tool called GINtool that can use regulon information to analyse transcriptome data. GINtool includes the option to take the activity mode of a regulator in account, which is important when a regulator can function both as an activator and repressor. GINtool also contains two novel graphical representations that greatly facilitate the visual inspection of regulon-based transcriptome analyses. Additional features of GINtool includes the evaluation of transcriptome data using functional categories, and the analysis of gene expression differences within operons. To ease the analyses and downstream processing of figures, GINtool has been developed as an add-in for Excel.

## INTRODUCTION

Genome-wide transcriptome analysis is a powerful tool to measure changes in gene regulation. However, due to the sheer number of genes, interpretation of the data necessitates a focus on a limited set of genes. Generally, this is accomplished by setting a threshold level for the fold-change and its statistical significance, often expressed as p-value. However, this may result in the loss of important information on gene regulation. Here we present a software tool, called GINtool, that analysis transcriptome data by using regulon information. This has several advantages. Firstly, by averaging the expression of all genes in a regulon the statistical relevance can be better assessed, secondly, the visual analysis of transcriptome data is simplified since the number of data points is reduced, and thirdly, such analysis provides a direct insight into the regulation networks responsible for the transcriptome effects.

The way transcriptome data is visualized has barely changed over the years, and is limited to Venn diagrams, Volcano plots, heat or tree maps (1). Generally, these representations show global effects and can be difficult to interpret. In GINtool, we provide two novel ways that facilitate the graphical visualization of changes in regulon activities or functional categories. Finally, GINtool also contains an option to visually inspect gene regulation within operons. This feature can be useful to check whether co-regulation of genes occurs. GINtool is provided as an Excel plugin so that graphs and tables can be analysed and adjusted using standard Excel tools. Here we illustrate the different capabilities of GINtool by analysing transcriptome data from the Gram-positive model bacterium *Bacillus subtilis* for which 223 regulons have been described, so far(2–4).

## MATERIAL AND METHODS

### Bacterial strains and general growth conditions

Bacterial strains and plasmids used in this study are listed in Supplementary Table S1. Nutrient Luria-Bertani medium (LB, containing 10 g/l tryptone, 5 g/l yeast Extract, 10 g/l NaCl) was used for general growth of both *B. subtilis* and *E. coli*. Supplements were added as required: kanamycin (50 μg/ml), spectinomycin (150 μg/ml), ampicillin (100 μg/ml), and IPTG (1 mM). For *B. subtilis* DNA transformation, the Spizizen-plus and Spizizen-starvation media (SMM, containing 15 mM (NH_4_)_2_SO_4_, 80 mM K_2_HPO_4_, 44 mM KH_2_PO_4_, 3 mM tri-sodium citrate, 0.5 % glucose, 6 mM MgSO_4_, 0.2 mg/ml tryptophan, 0.02 % casamino acids, and 0.00011 % ferric ammonium citrate ((NH_4_)_5_Fe(C_6_H_4_O_7_)_2_) were used and transformants were selected on LB-agar plates with antibiotic selection (5).

### Plasmid and strain construction

The effect of xylanase overproduction on the transcriptome was tested in *B. subtilis* strain BWB09 that lacked the native xylanase expressing gene *xynA* and amylase gene *amyE*. These genes were removed from the tryptophan-prototrophic (trp+) wildtype *B. subtilis* strain BSB1 (6) by means of a marker-free clean-deletion procedure (7). Briefly, purified DNA amplicons of *xynA*-upstream (primer pair BW45 & BW46), *xynA*-downstream (BW41 & BW44), *Sp*^*R*^*-mazF* cassette (BW05 & BW06) and *xynA* (BW47 & BW48) were fused by overlap PCR (BW45&BW44) to make the recombinant DNA construct. All primers used are listed in Supplementary Table S2. Subsequently, the construct was transformed directly into competent BSB1 cells and transformants were selected on spectinomycin selective plates. Next, colonies were inoculated in LB liquid containing 1 mM IPTG, which induced the MazF toxin expression, resulting in excision of the deletion cassette. Cells were again spread on LB agar to obtain single colonies. Cells from the edge of single colonies were restreaked on LB and spectinomycin plates, and those that grew on LB agar plates, but not on LB spectinomycin plates, were expected to have the *Sp*^*R*^*-mazF* cassette and *xynA* removed. The marker-less deletion was verified by PCR with *xynA*-internal and *xynA*-external primers (BW42 & BW43). Subsequently, the *amyE* gene was deleted using the same clean-knockout method, eventually resulting in strain BWB09 (*trpC*+, Δ*xynA*. Δ*amyE*).

For the overproduction of XynA, we used plasmid pCS58 which was based on the multicopy expression plasmid pUB110 containing *xynA* cloned downstream of the strong constitutive *amyQ* promoter (8, 9). As a negative control, we constructed an empty plasmid based on pCS58 from which the *xynA* ORF was removed by cyclizing using PCR primers BW34 and BW35 and self-ligation, resulting in plasmid pBW17.

### RNA extraction for RNA-seq

To examine the effect of xylanase overproduction on the transcriptome of *B. subtilis*, the BWB09 strains containing either pCS58 or pBW17 were grown in LB medium at 37 °C for approximately 6 h when the cells entered the stationary growth and the cultures reached an OD_600_ of approximately 4. The cultures were inoculated from overnight cultures, and 50 μg/ml kanamycin was present in the medium to maintain the plasmids. The experiment was repeated one more time to provide a biological replicate.

RNA extraction was based on the methods described in (10, 11). Briefly, 2 ml cells were collected by centrifugation (20,000 x rcf) at 4 °C for 1 min. Cell pellets were resuspended in 0.4 ml ice-cold growth medium and added to a screw cap tube containing 1.5 g glass beads (0.1 mm), 0.4 ml phenol chloroform/isoamyl alcohol (P/C/I) mixture (25:24:1) and 50 μl 10 % SDS, vortexed to mix, and flash frozen in liquid nitrogen. Cell disruption was achieved by bead beating (Precellys 24). After centrifugation, RNA in the upper aqueous phase was ethanol-precipitated, washed twice with 70 % ethanol, air dried and dissolved in water. DNA was removed by DNAseI (NEB) treatment. The pure total RNA was then extracted by a second round of P/C/I extraction, followed by ethanol-precipitation and 70 % ethanol washing, and finally dissolved in water and stored at -20 °C.

### RNA-seq

Prior to deep-sequencing, ribosomal RNA (rRNA) was removed using the MICROBExpress™ Bacterial mRNA Enrichment Kit (Thermo Fisher). Subsequently, the RNA-seq libraries were constructed using the NEBNext^®^ Ultra™ II Directional RNA Library Prep Kit for Illumina^®^ (New England Biolabs) and NEBNext® Multiplex Oligos for Illumina® (New England Biolabs), according to the manufacturer’s protocols. Sequencing was performed on an Illumina NextSeq 550 System using NextSeq 500/550 High Output v2.5 kit (75-bp read length), and the raw data were processed using the web-based platform Galaxy. We aimed at a sequencing depth of around 7 million reads/ library, and eventually gained 7.28 million reads/ library on average. *Trimmomatic* was used to trim adaptor sequence and filter bad reads (12). Trimmed reads were aligned to the *B. subtilis* reference genome (NC_000913) with *Bowtie2* (13). After mapping, aligned reads were counted by *FeatureCount* referred to the BSU locus_tag in the annotation, detecting 4406 genes, pseudogenes and 178 RNAs (14). *Deseq2* was used to determine differentially expressed features between samples (15). The fold-change, p-value, and functional description of all genes are listed in Supplementary Table S3.

### GINtool

GINtool was written in C# as an Excel plugin (VSTO) and runs on Windows versions of Excel. An extensive manual and starter package with a GINtool execution file and data test files are provided as Supplementary data file “Gintool start”.

## RESULTS AND DISCUSSION

### GINtool analysis using regulon information

The genome of *B. subtilis* comprises approximately 4800 genes and regulatory RNAs. So far 223 regulators have been characterised that regulate these genes and RNAs (2–4). Detailed information on these regulators and their regulons can be found in Subtiwiki, the main knowledge repository for *B. subtilis* (3). This database contains up-to-date and easy to use tables, listing gene functions, regulons, functional categories, operon structures and other information. To be able to analyse transcriptome data using these different datasets, we developed the Excel plugin GINtool. To illustrate the use of GINtool, we used original RNA-seq data from a *B. subtilis* strain overexpressing the industrial relevant xylanase XynA and compared this with RNA-seq data from a strain that does not express XynA (see Material and Methods for details).

A visual insightful way to show how regulons are affected is by plotting the genes of regulons according to their fold-change. Fig. 1 shows the result of such a spread plot made by GINtool using the XynA overproduction transcriptome data. The regulons are ranked according to the average fold-change of their related genes. GINtool also provides a table listing the average fold-change and p-value of regulons (see manual). Of note, the p-values do not follow a normal distribution so they first have to be transformed using a Fisher-Z transformation into z-values. The average of these z-values is then back-transformed resulting in the average p-value. To facilitate the visualization of the most important regulons, GINtool has the option to show only the most strongly affected regulons, based on either average fold-change or average p-value (Fig. 1, inset).

**Fig. 1.**
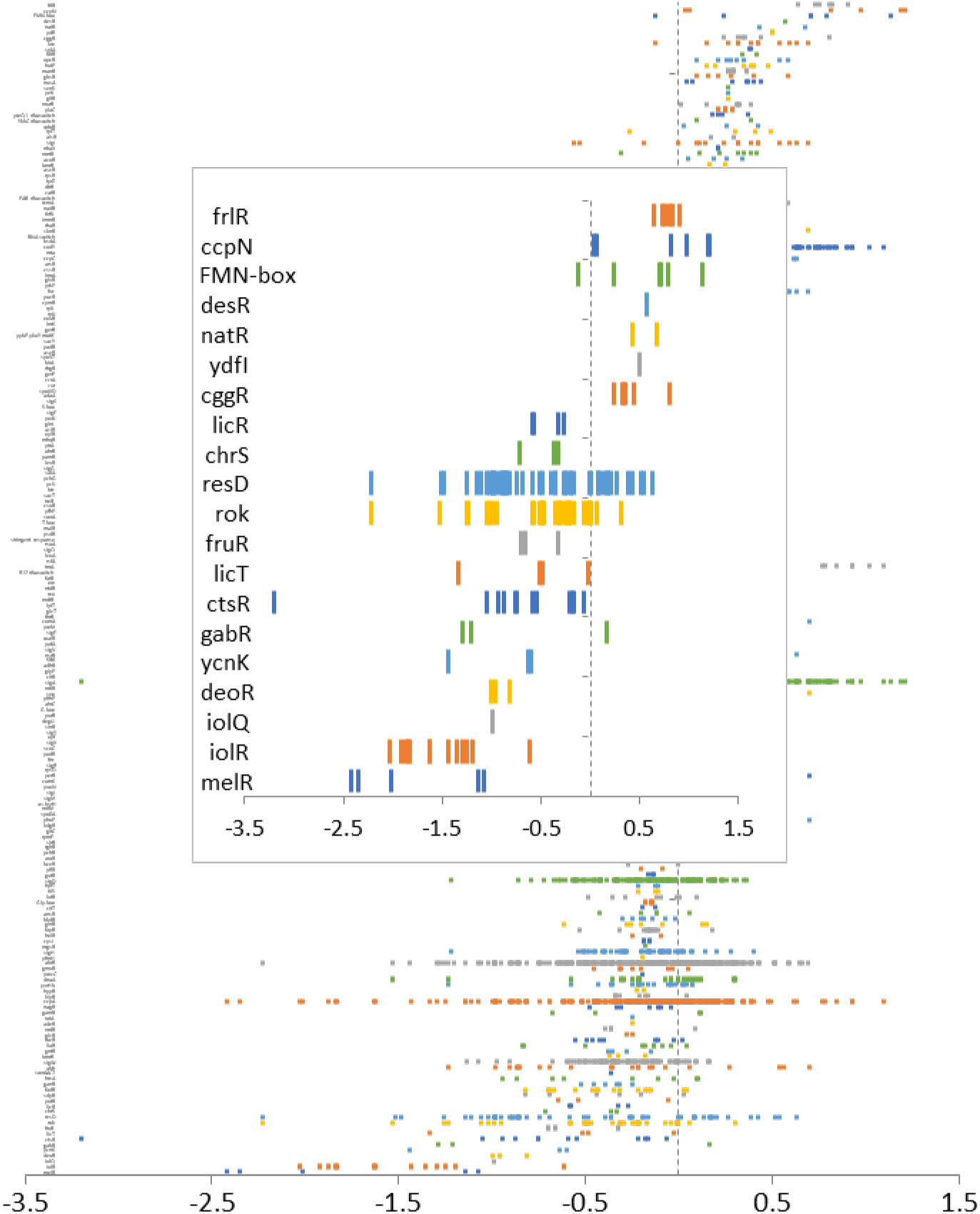
GINtool ranking of regulons based on average fold-change using spread plots. X-axes indicate fold-change values in log2 scale for our data. Each small bar represents the fold-change value of a gene, and each row represents all genes of a regulon. Regulons are ranked based on average fold-change in descending order. Names of regulons are listed at the left. The inset shows the option to display only the most up- and down regulated regulons (top 20, in this case).

### Visualisation using bubble plots

Due to the large number of regulons, it is difficult to determine from Fig. 1 which regulons are most relevant. Moreover, some regulons show a wide spread of fold-changes whereas others show a narrower range, suggesting a more uniform, thus robust regulation. To capture the latter, we calculated the median-absolute-deviation (MAD) of the fold-change values of regulon genes. The MAD value is a robust approximation of the spread around the median value (16). The median was chosen since it is less prone to outliers and returns the ‘middle’ value of the observed fold-changes after sorting. It emerged that these MAD values provide a useful means to visualize transcriptional differences between regulons when plotted against the average fold-change of regulons, as shown in Fig. 2A. In fact, we found that such a graph shows more details than plotting the average fold-changes against the average p-values, as in Volcano plots. The bubble plot layout was chosen so that the size of a regulon, *i*.*e*. the number of genes in a regulon, can be displayed as well, but in Excel this option can be easily discarded. Finally, the average p-values of regulons were also included by using different colour intensities. Excel can remove less relevant regulons, those with high p-values, to make a graph more clear, as is shown in Fig. 2B. In this case we also added the names of regulons. These labels can be easily added, moved or removed in Excel.

**Fig. 2.**
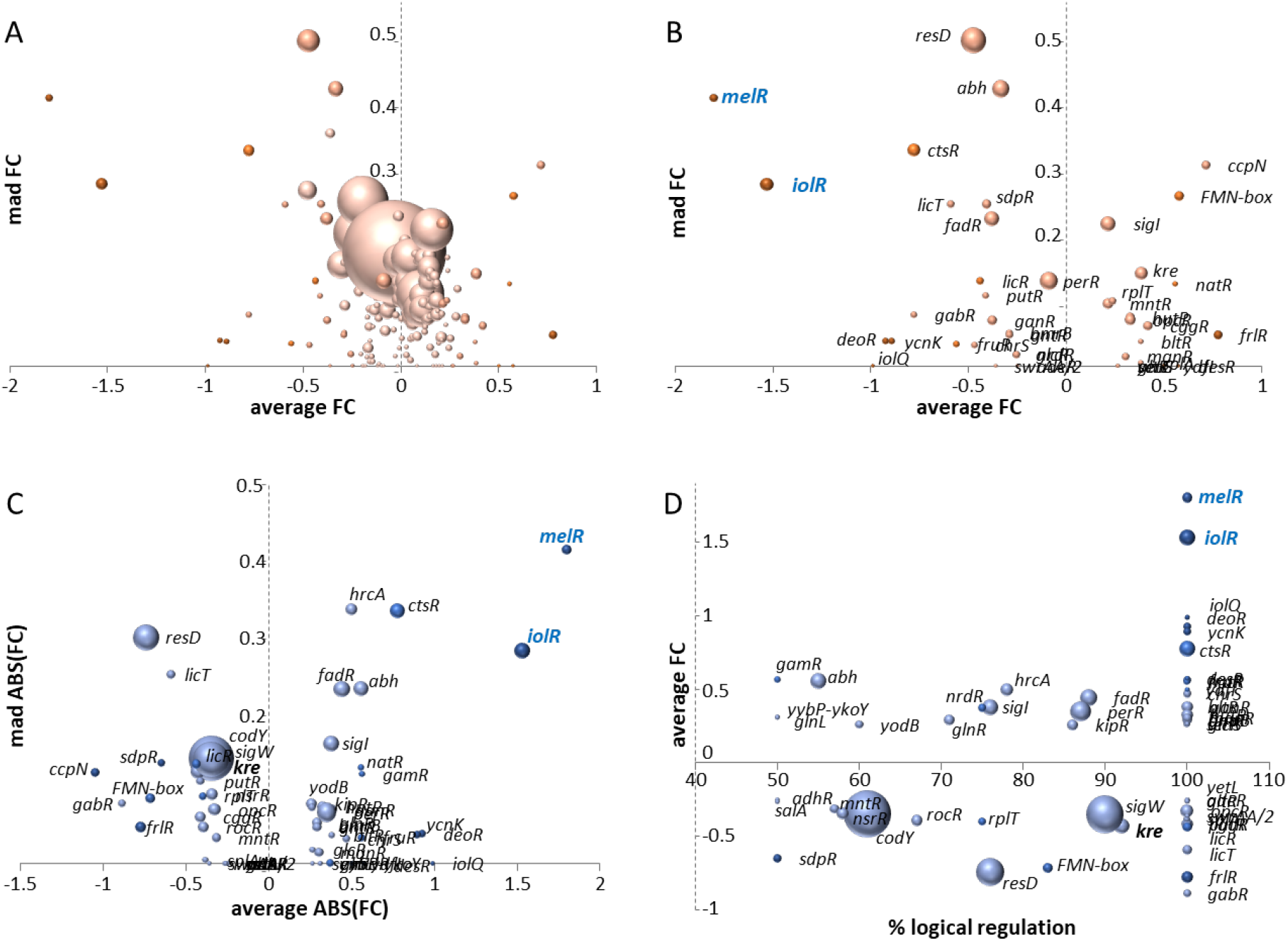
GINtool visualization of regulon data by means of bubble plots. (A,B) Regulon bubble plots showing average fold-change (FC, log2 scale) of regulons plotted against the Mean Absolute Deviation (MAD) of the fold-change (mad FC, log2 scale). The size of bubbles corresponds to the number of genes in a regulon. The colour intensity of the bubbles indicates the average p-values of the genes in a regulon, and goes from dark to light according to average p-values of <0.0625, <0.125, <0.25, <0.5 and >=0.5. Panel (A) shows information on all regulons, while panel (B) shows the regulons with average p-values <0.5. (C) Regulon bubble plots corrected for regulation activity (directionality). Negative or positive average fold-change values indicate that the activity of the corresponding regulator is inhibited or activated, respectively. As a consequence, regulons like IolR and MelR (marked in blue) switch position in plot C compared to plot B. (D) Average fold-change from C plotted against the fraction of regulon genes used to calculate the average fold-change. For clarity, only regulons with average p-values <0.5 are shown in B-D.

### Using directionality of regulation

Many transcription factors can function both as activator and repressor. For example, the *B. subtilis* response regulator Spo0A regulates 143 genes of which approximately half by activation and the other half by repression. As a consequence, the Spo0A regulon will show an average fold-change approaching zero, no matter whether Spo0A is active or not. To overcome this problem, GINtool has the option to take into account the directionality of regulation, i.e. whether a gene is activated or repressed by a regulator. This information can be found in the regulon file from the Subtiwiki website. Of note, sigma factors were considered positive regulators in this analysis. GINtool can then calculate the average fold-change, related MAD and average p-values for two scenarios, using either the number of genes whose fold-change direction (negative/positive) fitted with an activated regulator or with a repressed regulator. The activity that is in agreement with the fold-change direction of most regulon genes is then considered the most likely activity of the regulon. This information can also be plotted as a bubble graph, as shown in Fig. 2C. In this case the average fold-change, MAD and p-values are based only on the genes that show a regulation related to the most likely activity mode of the regulator. What is noticeable is that the average fold-change direction differs for several regulons. E.g. MelR and IolR show a negative fold-change in Fig. 2B but a positive fold-change in Fig. 2C. The reason for this is that both MelR and IolR function as repressors, so a downregulation of the genes that they regulate, indicated in Fig. 2B, means that both regulators are active, therefore GINtool assigns them a positive value in Fig. 2C. This assignment is important when dealing with regulators like Spo0A, for which the average fold-change is made up of both up- and down-regulated genes. The information in the bubble plots is also provided in tables (see manual), enabling customization of the regulon data display.

### Including fraction of regulated genes

The graph in Fig. 2C presents, by means of bubble size, the number of genes that fit the most likely activity direction (repression/activation) of a regulon. However, this number does not indicate what fraction of regulon genes show this mode of activity. For example, when 51 % of the genes of a regulon show a regulation that corresponds to the regulator being activated, then this regulation is not very relevant, considering that 49 % of the regulon genes shows regulation that corresponds with a repressed regulator. Yet, for large regulons 51 % will still show a considerable bubble size. GINtool can work around this problem by plotting the average fold-changes against these calculated fractions, as shown in Fig. 2D. For example, this graph strongly suggests that the activity of the regulator Kre is slightly repressed, whereas this information would have been lost when a fold-change cut-off of 2 would have been applied. GINtool provides this information also in a table format (see manual).

### Regulon regulation information per gene

When listing the most up- or down-regulated genes in a table, it can be useful to also indicate the regulator that might be responsible for the observed regulatory effect. GINtool can link the information shown in Fig. 2D, i.e. average fold-change and best fitting regulon fraction, to individual genes in a table (Fig. 3). It will show this data for all the regulons to which the gene belongs and leaves the interpretation which regulator is most likely up to the user.

**Fig. 3.**
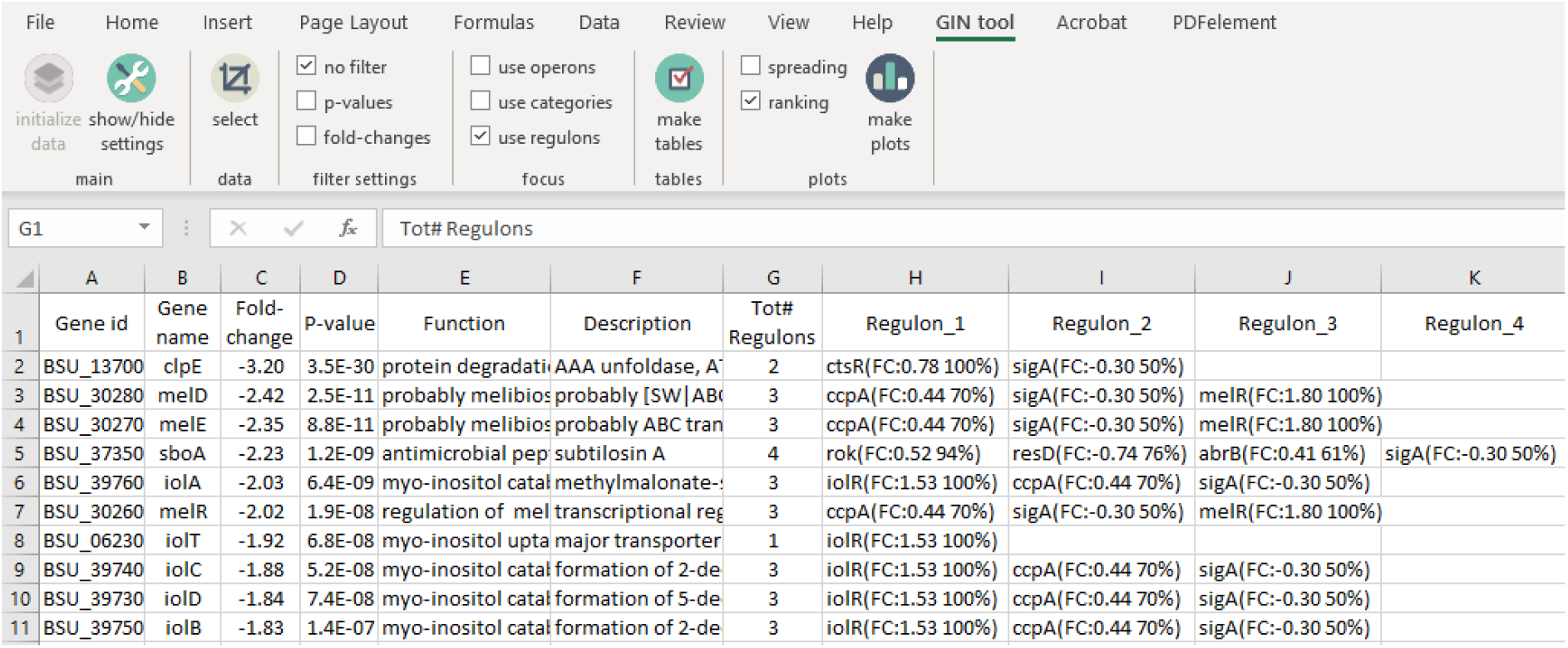
Linking regulators to genes in GINtool. Example of a GINtool output table showing the main transcriptome information of genes with the regulons to which a gene belongs. Column G lists the number of regulons to which genes belong. Columns H, I, etc. list these regulons with their most likely activity (negative for repression and positive for activation), the average fold-change (FC) for the most likely activity, and the fraction (%) of genes of the regulon that determines the most likely activity, respectively.

### Expression differences in operons

Genes in an operon should show the same regulation. However, due to (arbitrary) fold-change and/or p-value cut-offs, it often seems as if only a few genes of an operon are significantly regulated. GINtool gives the opportunity to visually examine expression differences of genes in operons. For example, Fig. 4 shows a discrepancy in the *yxaJ* operon, with the first gene being upregulated whereas the second gene is clearly downregulated. This suggests that the second gene in the operon (*yxaL*) is regulated by an unknown regulator and that this operon is not appropriately annotated. Indeed, an extensive transcriptomic study (6), which is accessible in the Expression Browser of Subtiwiki, suggests that *yxaJ* and *yxaL* are, at least partially, regulated in different ways. In fact, there is a 101 bp spacer region between both genes, thus ample space for a promoter to regulate *yxaL*.

**Fig. 4.**
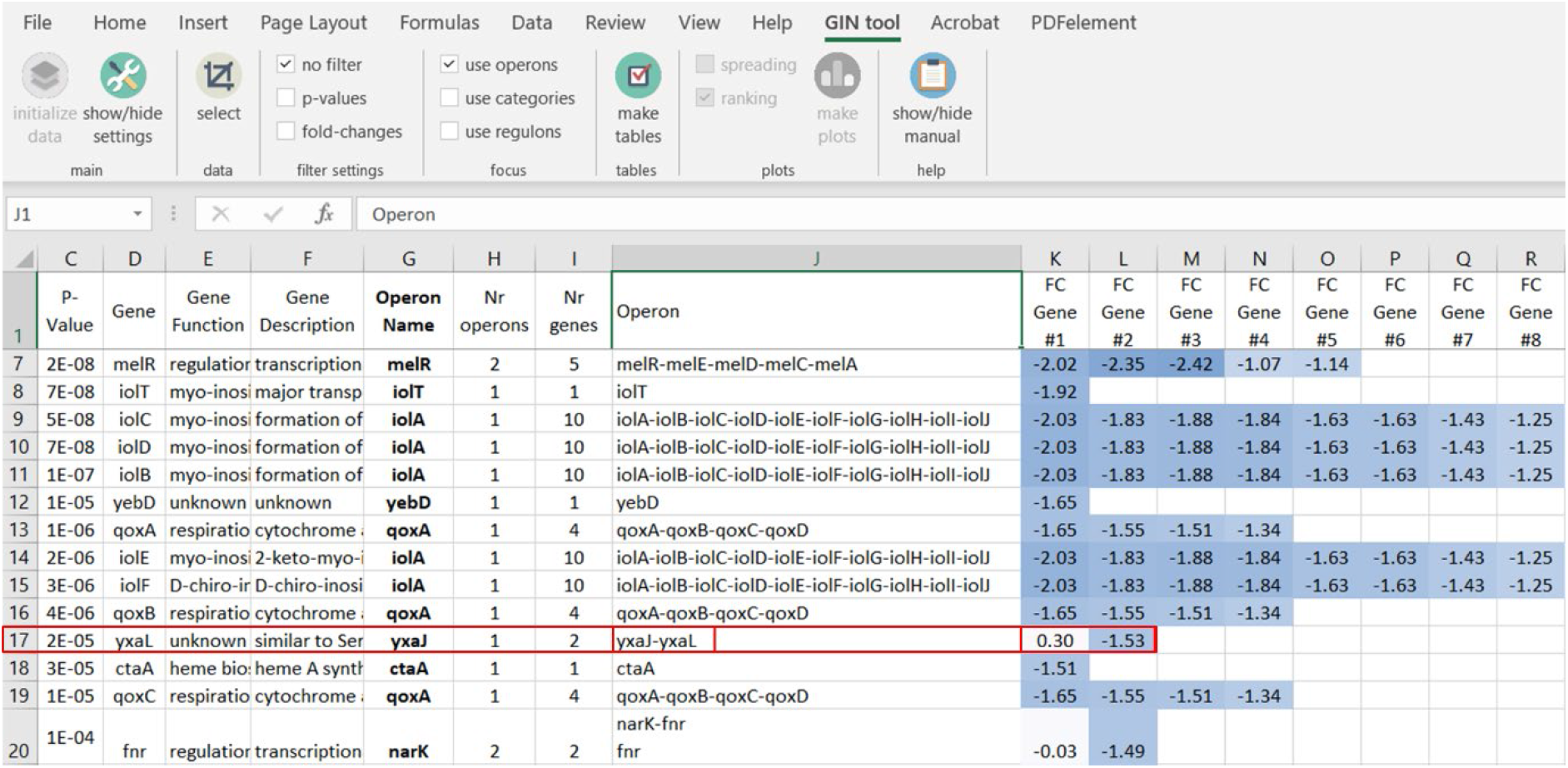
Expression differences of genes in operons in GINtool. Example of a GINtool output table showing the main transcriptome information of genes, including the operon(s) the gene belongs to, with the fold-changes of the genes in these operon(s). Column H indicates the number of operons a gene belongs to, column J shows the gene context of these operons, and the rest of the columns (K, L, etc.) show the expression fold-change of these genes. Fold-changes were colour coded using the conditional formatting function of Excel. The red box shows an example of an operon (yx*aJ*-*yxaL*) with an illogical fold-change distribution.

### Use of functional categories

*B. subtilis* is one of the best studied bacteria, but for most bacterial species their gene regulation networks are not so well characterized. In these cases, the use of functional categories is a good option to analyse transcriptome data, since during the annotation of genomes, genes are classified in functional categories. For *B. subtilis* more than 350 functional categories and sub-categories have been defined. GINtool also can make spread plots and bubble plots, as those shown in Fig. 1 and 2, using functional category information (see manual).

### Conclusion

Here we describe a new software tool to analyse transcriptome data using regulon and functional category information, and present the results in novel, easy to inspect, graphical formats. As far as we know, GINtool is the first analysis tool that takes regulation directionality into account, which is crucial to analyse the effect of regulators that work both as activator as well as repressor.

## Supporting information

GINtool starter package

## DATA AVAILABILITY

RNA-seq data have been submitted to and are accessible in the Gene Expression Omnibus (GEO), accession number GSE208571. The GINtool source and executable can be downloaded from GitHub (ScienceParkStudyGroup/GINtool: Gene Information Network add-in for Excel).

## ACKNOWLEDGEMENTS

We would like to thank Yoena Nossent, Niels Huiberts, Pascal Maas and Anne van Winzum for coming up with the GINtool name. BW and MK were supported by a NWO-TTW (17833) grant awarded to JL and LWH.

