## Supplementary material for "Transcriptome analyses using regulon, functional category, and operon information with GINtool": Table S1 and Table S2

**Table S1. Strains and plasmids used in this study.**

| Strain | Genome type | Source |
|---|---|---|
| BSB1 | *B. subtilis wildtype* 168 trp+; | Lab stock |
| BWB09 | BSB1 trp+; Δ*xynA*, Δ*amyE* | This study |
| **Plasmid** | | |
| pCS58 | $P_{amyQ}$-*xynA, bleo(Km)* | Lab stock |
| pBW17 | $P_{amyQ}$-*empty, bleo(Km)* | This study |

**Table S2. Primer sequences used in this study.**

| Name | Sequence (5'-3') | Target |
|---|---|---|
| BW05 | CTAATTGAGAGAAGTTTCTATAGAATTTT | SpR-mazF-Fw |
| BW06 | CTACCCAATCAGTACGTTAATTT | SpR-mazF-Rv |
| BW34 | AAAGGAGCGATTTACATATGTAACAGATCATCCTTAATCA | pEmpty1-Fw |
| BW35 | TGATTAAGGATGATCTGTTACATATGTAAATCGCTCCTTT | pEmpty1-Rv |
| BW41 | CAGATCATCCTTAATCAGGGGTAGCTAACG | XnyADn-Fw |
| BW42 | GAAACTTCTCTCAATTAGATTTCATGTAAACCGAGAACCA | XnyAEx-Rv |
| BW43 | GTTATAACGCACCTTCCATT | XynAIn-Fw |
| BW44 | AACTTTTTAACTAGAAAGCGCACGTAGTGTGATTATATCACCGTG | nyADn-Rv |
| BW45 | GCAAAAGCCCTTATGAGGGCTTTTTTAATTGTTGTTTGCAGTAAC | XnyAUp-Fw |
| BW46 | ACCCCTGATTAAGGATGATCTGATGTTACCTCCTATAATATTTTTTCCG | XnyAUp-Rv |
| BW47 | AATTAACGTACTGATTGGGTAGTTCTTAGTTGGATTATCGGCAGC | XnyA-Fw |
| BW48 | GGATGATCTGTTACCACACTGTTACGTTAGAACTTCCACTAC | XnyA-Rv |
| BW49 | ATGATCAATTGGGGGCCGTTTTAACGATTGCTGCC | AmyEUp-Fw |
| BW50 | TCCCGTCTAGCCTTGCCCTCTTGACACTCCTTATTTGA | AmyEUp-Rv |
| BW51 | GGGCAAGGCTAGACGGGACTTACCGAAAGAAA | AmyEDn-Fw |
| BW52 | TATAGAAACTTCTCTCAATTAGCCCGCTCTTTTGGCAGGCCGC | AmyEDn-Rv |
| BW53 | AACGTACTGATTGGGTAGGCCATTCAGACATCTCCGA | AmyE-Fw |
| BW54 | CAGACCTGGCATTGATCGTGCCTGTCAGTTTAC | AmyE-Rv |
| BW135 | CTTTGAAGCTTGGCTGGTC | AmyEEx-Fw |
| ZT080 | CACATTGTGAAATCTATTGACCGCAGTG | AmyEIn-Rv |
