## Supplementary material for "Transcriptome analyses using regulon, functional category, and operon information with GINtool": GINtool starter package: GIN tool manual.pdf

### GINtool manual

Version: 19 October 2022

**Contents**


1. Introduction to the tool 2

2. Installation of GINtool 2

3. Overview of the tool 2

4. Preparing GINtool for analyses 6

5. Analysis of operons 8

6. Analysis of regulons 10

7. Analysis of functional categories 22

8. Overview of output sheets 34

9. Other ways to use GINtool data 36

10. Calculations of MAD & p values 36

#### 1. Introduction to the tool

GINtool is an Excel plugin that is used for a comprehensive analysis of transcriptome data based on information on operons, regulons and functional categories.

This manual describes how to use GINtool. For each type of analysis, examples are shown.

#### 2. Installation of GINtool

##### 2.1. Preparation

Before installing GINtool, close Excel.

##### 2.2. Installation

Click setup.exe to install GINtool.

> When clicking on setup.exe the following error message might appear: "*Customized functionality in this application will not work because the certificate used to sign the deployment manifest for GINtool or its location is not trusted. Contact your administrator for further assistance.*"
>
> In this case do the following:
>
> 1) Right click on setup.exe;
>
> 2) Select properties then the Digital Signatures tab;
>
> 3) Select "WIN-SMURF\frans" then click Details;
>
> 4) Click View Certificate and Click Install Certificate, go to Current user then Details and choose "Place all certificates in the following store";
>
> 5) Browse and select "Trusted Root Certification Authorities Store". Finally, finish and close the windows;
>
> 6) Click setup.exe again to install GINtool.

**!** In case Excel at some point starts to behave strangely, then uninstall GINtool and reinstall GINtool.

#### 3. Overview of the tool

##### 3.1 GINtool interface

GINtool is an Excel plugin that is added as a tab to the Excel toolbar after installation (see section 2). Figure 1 shows an overview of the tool. After clicking on the GINtool tab (Figure 1A, red box), the analysis bar appears (Figure 1B). From here you can move to the settings (main), select your data (select), activate filters (filter settings), choose the focus of your analysis (focus), create tables (tables), create plots (plots) and find this manual (help).

Via "show/hide settings" (Figure 1B, main) the settings bar appears (Figure 1C). The reference files on gene info, operons, regulons, categories and regulon info can be uploaded (reference files), cut-offs can be determined (cut-offs) and sorting directions of the results can be chosen (sort direction). The columns of the reference files (see section 3.2) are linked to the tool by clicking on the reference file buttons and selecting the appropriate column names (column mapping, *e.g.*, for regulons in Figure 1D, see section 4.1).

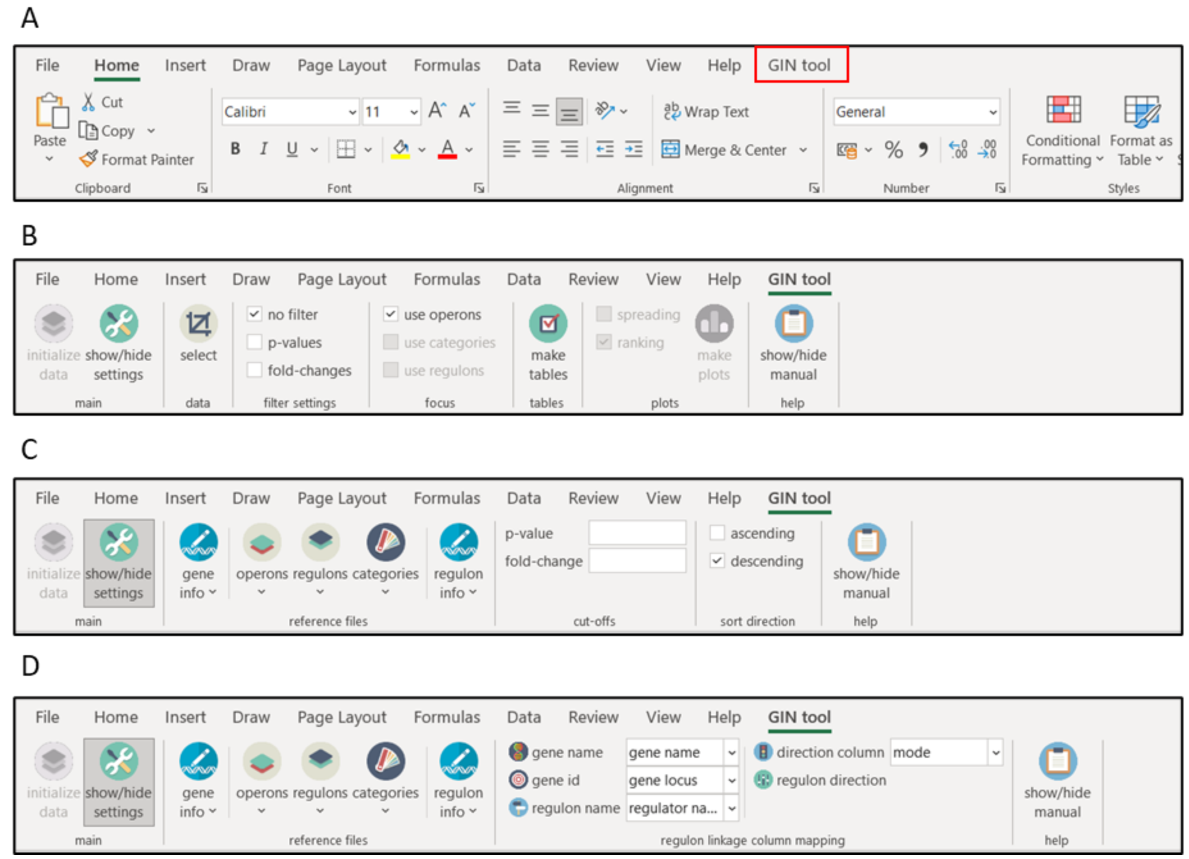


Figure 1: Overview of the GINtool plug-in. A) Location of GINtool tab in toolbar; B) Analysis bar; C) Settings bar; D) Regulon column mapping.

#### <u>3.2 Overview of input reference files</u>

GINtool works with 5 different information files: (i) A file containing information on the function of genes ("<u>gene info</u>"), (ii) A file indicating how genes are located in operons ("<u>operons</u>"), (iii) A file indicating to which regulon the gene belongs ("<u>regulons</u>"), (iv) A file indicating to which functional category the gene belongs ("<u>categories</u>"), and (v) A file containing information on the function of regulons ("<u>regulon info</u>"). These files can be imported either as txt files (*.csv) or Excel files (*.xlsx). The genes are linked to the information by either gene name or unique identifiers (ID).

The files shown in the examples are based on information obtained from *Subtiwiki*, a database for the model bacterium *Bacillus subtilis*. Figure 2 shows an overview of the different (.csv) reference files. Each reference file starts with a header for each of the columns in the file.

- The gene info reference file contains four columns: locus_tag, gene, function and description (Figure 2A). The gene ID, the gene name, gene function, and functional description are separated by commas.

- The operon reference file contains one column: genes (Figure 2B). The operons are indicated as a list of gene names that are connected (e.g., yokI-yokJ-yokK-yokL). Single locus genes are indicated by single gene names (e.g., ywqG).
- The regulator reference file contains five columns: regulator locus, regulator name, mode (type of regulation), gene locus and gene name, separated by commas (Figure 2C). Importantly, GINtool does not use the regulator ID (*i.e.*, regulator locus), because the regulator name is not always defined by a single regulator protein, for example the "stringent response" regulon.
- The category reference file contains three columns: category id, category and gene locus (Figure 2D). There are several layers of subcategories, and in total a depth of 5 categories with subcategories indicated by the identifier SW.x.x.x.x.x for the final subcategory. The category ID is followed by the description of the category/subcategory, and then the gene ID, separated by commas.
- The regulon info reference file contains three columns: regulon, regulon size (nr of genes) and function (Figure 2E). This information is not available in *Subtiwiki*, but was based on the "gene info" information, where the function of the gene encoding the regulator is described. The information required is the name of the regulon, the number of genes in a regulon and the function of the regulon, separated by commas.

**!** The headers of the columns can be adjusted. When uploading the files to GINtool, the header names will be mapped to the corresponding elements (see section 4.1).

**!** GINtool only works when the "gene info column mapping" has been completed. It is also crucial to properly fill in the "regulon linkage column mapping", "category linkage column mappings" and/or "regulon info column mapping", when used.

**!** Gene info is mandatory for each analysis, in combination with at least the reference file(s) for either operons, regulons or categories.

A

| | A | B | C | D | E | F | G |
|---|---|---|---|---|---|---|---|
| 1 | locus_tag,gene,function,description | | | | | | |
| 2 | BSU_13700,clpE,protein degradation,"AAA unfoldase, ATPase subunit of the ( | | | | | | |
| 3 | BSU_30280,amyD,probably melibiose uptake,probably [SW\|ABC transporter] | | | | | | |
| 4 | BSU_30300,melA,melibiose utilization,alpha-galactosidase | | | | | | |
| 5 | BSU_30290,amyC,probably melibiose uptake,probably [SW\|ABC transporter] | | | | | | |
| 6 | BSU_30270,msmE,probably melibiose uptake,probably ABC transporter for m | | | | | | |
| 7 | BSU_37350,sboA,antimicrobial peptide,subtilosin A | | | | | | |
| 8 | BSU_30260,msmR,regulation of melibiose utilization,transcriptional regulato | | | | | | |
| 9 | BSU_39760,iolA,myo-inositol catabolism,methylmalonate-semialdehyde deh | | | | | | |
| 10 | BSU_06230,iolT,myo-inositol uptake,major transporter for inositol | | | | | | |

B

| | A | B | C | D | E | F | G |
|---|---|---|---|---|---|---|---|
| 1 | genes | | | | | | |
| 2 | ywqG | | | | | | |
| 3 | yokI-yokJ-yokK-yokL | | | | | | |
| 4 | mbl | | | | | | |
| 5 | coaX-yacC-yacD-cysK | | | | | | |
| 6 | htrC | | | | | | |
| 7 | rpmGA | | | | | | |
| 8 | scoC | | | | | | |
| 9 | pepF | | | | | | |
| 10 | pucJ-pucK-pucL-pucM | | | | | | |

C

| | A | B | C | D | E | F | G |
|---|---|---|---|---|---|---|---|
| 1 | regulator locus,regulator name,mode,gene locus,gene name | | | | | | |
| 2 | BSU_36420,spoIIID,activation,BSU_06900,cotJB | | | | | | |
| 3 | BSU_36420,spoIIID,activation,BSU_06890,cotJA | | | | | | |
| 4 | BSU_36420,spoIIID,activation,BSU_06910,cotJC | | | | | | |
| 5 | BSU_36420,spoIIID,activation,BSU_06920,yesJ | | | | | | |
| 6 | BSU_36420,spoIIID,activation,BSU_06930,yesK | | | | | | |
| 7 | BSU_15320,sigE,sigma factor,BSU_06900,cotJB | | | | | | |
| 8 | BSU_15320,sigE,sigma factor,BSU_06890,cotJA | | | | | | |
| 9 | BSU_15320,sigE,sigma factor,BSU_06910,cotJC | | | | | | |
| 10 | BSU_15320,sigE,sigma factor,BSU_06920,yesJ | | | | | | |

D

| | A | B | C | D | E | F | G |
|---|---|---|---|---|---|---|---|
| 1 | category id,category,gene locus | | | | | | |
| 2 | SW.6.9.1,6S RNA,BSU_misc_RNA_32, | | | | | | |
| 3 | SW.6.9.1,6S RNA,BSU_misc_RNA_41, | | | | | | |
| 4 | SW.3.1.5.2,A/P endonucleases,BSU_40880, | | | | | | |
| 5 | SW.3.1.5.2,A/P endonucleases,BSU_25130, | | | | | | |
| 6 | SW.3.1.5.2,A/P endonucleases,BSU_22340, | | | | | | |
| 7 | SW.2.6.5.4,ABC transporters for the uptake of iron/ siderophores,BSU_3329( | | | | | | |
| 8 | SW.2.6.5.4,ABC transporters for the uptake of iron/ siderophores,BSU_3331( | | | | | | |
| 9 | SW.2.6.5.4,ABC transporters for the uptake of iron/ siderophores,BSU_0163( | | | | | | |
| 10 | SW.2.6.5.4,ABC transporters for the uptake of iron/ siderophores,BSU_0162( | | | | | | |

E

| | A | B | C | D | E | F | G |
|---|---|---|---|---|---|---|---|
| 1 | regulon,regulon size (nr of genes),function | | | | | | |
| 2 | Abh,22,regulation of gene expression during the transition from growth to sta | | | | | | |
| 3 | a-box,1,"termination/antitermination pbuE (putative hypoxanthine exporter, | | | | | | |
| 4 | AbrB,271,regulation of gene expression during the transition from growth to : | | | | | | |
| 5 | AcoR,4,regulation of acetoin utilization | | | | | | |
| 6 | AdaA,3,adaptive response to alkylative DNA damage | | | | | | |
| 7 | AdeR,1,control of L-alanine dehydrogenase expression (alanine utilization) | | | | | | |
| 8 | AdhR,2,regulation of the protective response to formaldehyde and methylgly | | | | | | |
| 9 | AhrC,16,transcriptional regulator of arginine metabolic genes | | | | | | |
| 10 | AlsR,2,regulation of acetoin synthesis | | | | | | |

Figure 2: Overview of input files. A) Gene info; B) Operons; C) Regulons; D) Categories; E) Regulon info.

#### 3.3 Overview of input dataset

Datasets that are analyzed with GINtool are the result of a differential expression analysis of your transcriptomics data. The file includes a column with gene names, fold-changes and p-values (or a related statistical classifier). Figure 3 shows an example. The fold-changes can be log2 transformed, but do not have to. Importantly, a minus sign (-) is required to indicate downregulation. Remove non-numerical values (like “NA”) for fold-change and p-values.

| | A | B | C | D |
|---|---|---|---|---|
| 1 | **locus** | **log2FC** | **p-val** | |
| 2 | BSU_11549 | 2.002516245 | 0.096155 | |
| 3 | BSU_21030 | 1.62902704 | 0.22792 | |
| 4 | BSU_14629 | 1.601961295 | 0.002607 | |
| 5 | BSU_36720 | 1.504901617 | 0.152471 | |
| 6 | BSU_20300 | 1.504410801 | 0.181841 | |
| 7 | BSU_06319 | 1.503871939 | 0.177237 | |
| 8 | BSU_27809 | 1.476434051 | 0.218652 | |
| 9 | BSU_06120 | 1.440394618 | 0.039961 | |
| 10 | BSU_20620 | 1.409932161 | 0.183179 | |
| 11 | BSU_32090 | 1.395148599 | 0.019007 | |
| 12 | BSU_07190 | 1.392904273 | 0.180525 | |

Figure 3: Example of transcriptomics data. The data contains three columns: a column containing gene IDs, a column with log2-transformed fold-changes and a column with p-values.

**!** Gene IDs in the dataset need to match the gene IDs in the reference files. For example, BSU11549 is not the same as BSU_11549.

**4. Preparing GINtool for analyses**

*First prepare the transcriptomics data and reference files according to **section 3**.*

4.1 First use: upload files

Open your transcriptomics data in Excel and open GINtool via the tab in the toolbar (Figure 1A). Click "show/hide settings" to open the settings. Upload the reference files for gene info and the analysis you want to perform (*i.e.*, operon, regulon or functional categories) by clicking on the appropriate buttons and selecting the files (Figure 4A). After uploading each file, link the column names to the correct element ("show/hide column mapping", Figure 1D).

In case of regulon information, also determine regulon directions (Figure 4B). The left box called "undefined" lists all definitions found under the "direction column" which is labelled "mode" in the example. For a complete regulon analysis it is necessary to separate the different types of regulation into activation and repression by moving them either into the "up-regulated" or "down-regulated" box. When the type of regulation is unknown it can be left in the "undefined" box. Of note, we consider "sigma factors" activators. This information will be maintained in Excel for use in next sessions, as long as you use the same regulon file (see 4.3).

**!** Gene info is mandatory for each analysis, in combination with at least the reference files for either operons, regulons or categories.

4.2 Select transcriptomics data

After uploading and mapping the reference data, hide the settings by clicking "show/hide settings". The "select" button should now be enabled. If not, click "initialize data". Click the "select" button, select the appropriate columns of your transcriptomics data (Figure 4C) and click “OK”. The analysis bar will now be enabled.

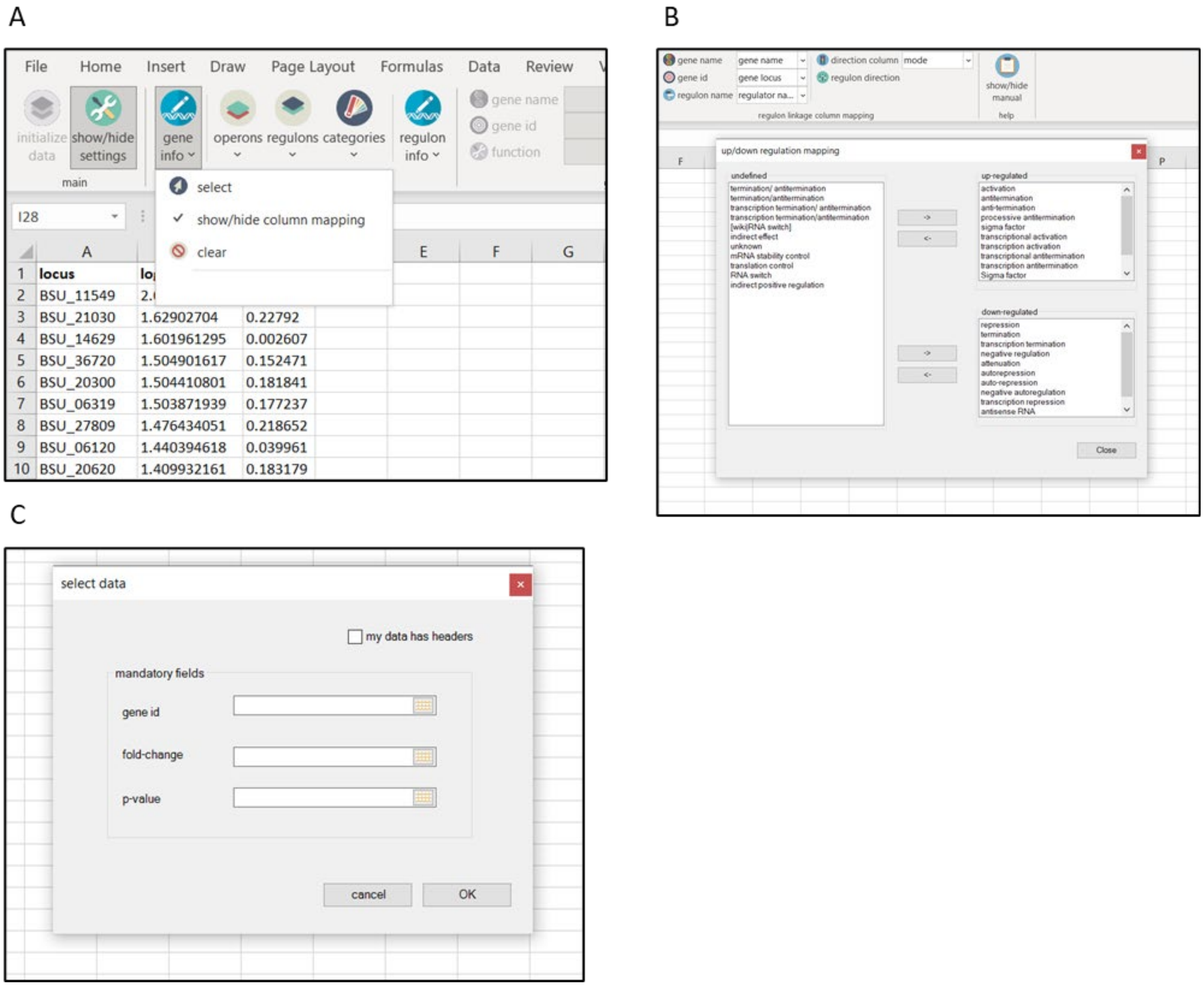


Figure 4: Uploading reference files and selecting your data. A) Selection of reference file for gene info; B) Selection of Up/down regulation mapping of regulons; C) Selecting of columns of transcriptomics data.

#### <u>4.3 Subsequent use: continue using GINtool after closing Excel</u>

After restarting Excel, you may need to repeat some of the steps described in 4.1 and 4.2. If the analysis buttons are disabled, you need to select the data columns again. If this is not possible, check if the reference files are still coupled by clicking on the arrow buttons beneath the reference file buttons. If the files are still coupled, click “<u>initialize data</u>” button in order to reload the files. If the files are not coupled, upload the files again, map the correct columns and click “<u>initialize data</u>”. Now you are again able to select your data and continue your analyses.

**5. Analysis of operons**

*First follow the steps described in* ***section 4****.*

5.1 Fold-changes genes in operons

To calculate fold-changes in operons, tick "use operons" in the "focus" window (Figure 5A, red box) and click "make tables" in the "tables" window. This will create a new sheet called "Mapped" (Figure 5B). In "Mapped", each row represents the information for one gene. Of note, the fold-changes in operon data begins in column K. To make this more clear, we used the Excel option "Conditional formatting" to highlight fold-change differences using colors. Column D shows the gene and column G the operon to which the gene belongs. The structure of the operon is indicated in column J, and column I shows the number of genes in the operon. Figure 5B, row 234, gives a clear example of an operon that is not regulated like an operon, since the two genes in the *yxaJ* operon shows very different fold-changes. Sometimes there are different transcriptional start sites in an operon, *e.g.*, this is shown in row 233 and 238 in Figure 5B. In this case column H gives the value 2 for the number of operons. Cell K236 is empty, because for some reason there is no proper fold-change data for the gene *yfnH*.

In the example above, the tick box "no filter" in the window "filter settings" is activated (Figure 5A), thus all transcriptomics data is used in this analysis. If you wish to prefilter genes, you can specify cut-off values for either the fold-change or p-value of the genes that are included in the analysis. Click "show/hide settings" to open the settings bar. The desired cut-offs can be determined in the cut-offs window (Figure 5C, red box). By default, no cut-offs are activated. To use a cut-off, click "show/hide settings" and tick "fold-changes" in the "filter settings" window in the analysis bar (Figure 5D, red box). The table can be created as described above. For example, the FC cut-off can be changed to 2. After filtering the genes based on a fold-change cut-off of 2, only 32 operons remain (Figure 5E). The same procedure can be followed to select genes based on p-values, or to select based on both cut-offs, by ticking the appropriate boxes in "filter settings" (Figure 5D).

**!** If the fold-changes in the data set are log2 transformed, then the FC cut-off is also interpreted as such.

**!** The fold-change cut-offs are absolute values, *i.e.*, fold 2 can be both 2-fold downregulation and 2-fold upregulation.

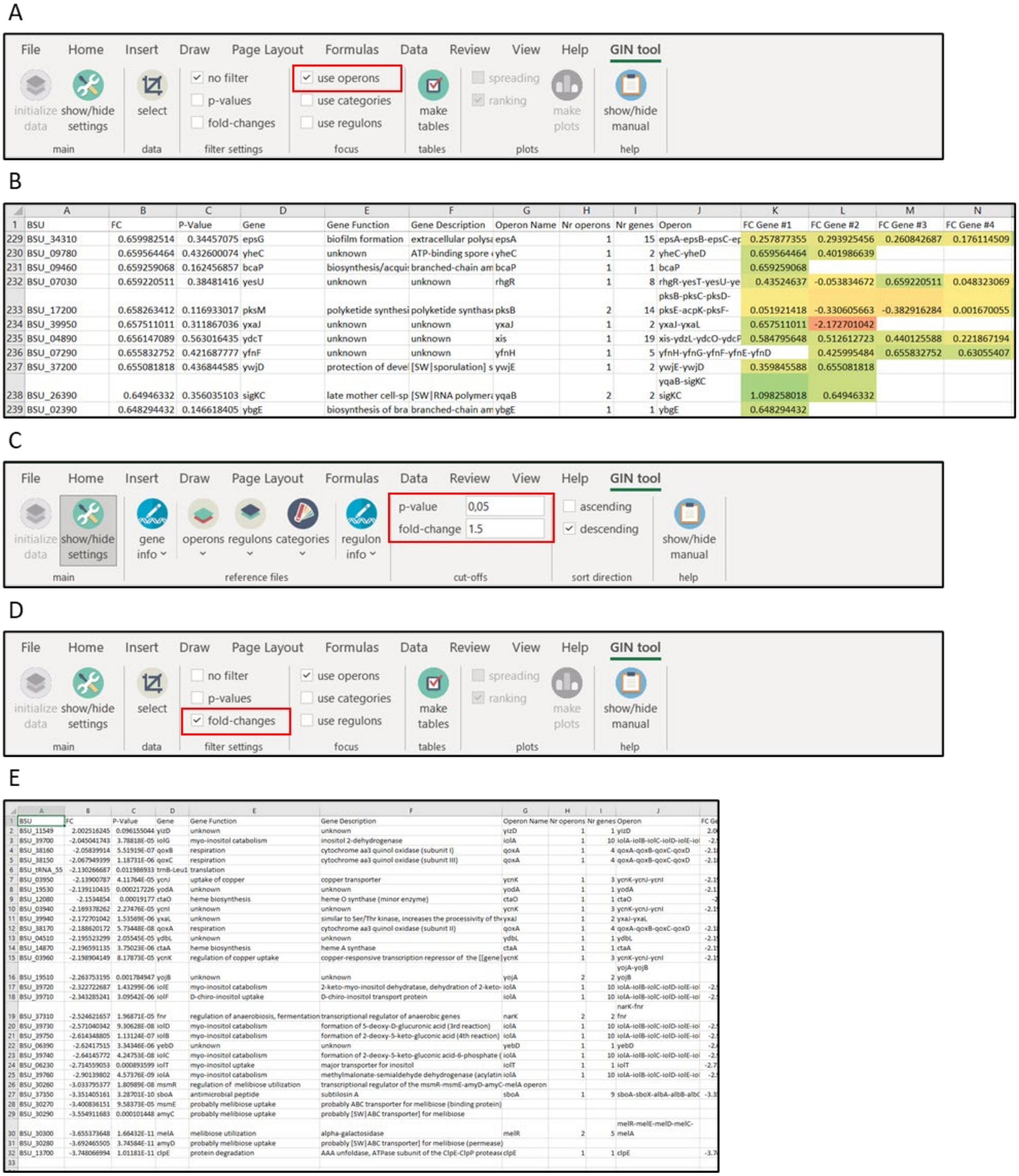


Figure 5: Analysis of operons. A) Selection of operons as focus of the analysis; B) Snapshot of "Mapped". Fold-changes are colored by conditional formatting; C) Cut-off settings; D) Activating fold-change filter; E) Overview of "Mapped" after filtering based on a fold-change of 2.

#### 6. Analysis of regulons

*First follow the steps described in **section 4**.*

6.1 Visual analysis of regulons using spread plots

GINtool gives the option to rank regulons based on the average fold-change of all genes in a regulon. The distribution of fold-changes of genes of regulons can be visualized in a spread plot. Start by ticking "use regulons" as "focus" and "spreading" for "plots" (Figure 6A). Click "make plots", the "Select categories/regulons" window will pop up (Figure 6B).

By clicking on "Regulons" in the left window "Available categories/regulons", all regulons become visible. Regulons can be selected and moved to the right window "Selection" by clicking on the single arrow button. It is also possible to select all regulons by clicking the double arrow. To produce the spread plot, click "Ok". This will create a new sheet called "RegSpreadPlot", displaying the fold-changes of all genes in the selected regulons. The regulons are ranked based on the average fold-change (Figure 6C). The x-axis indicates the fold-change. In the example all regulons were selected.

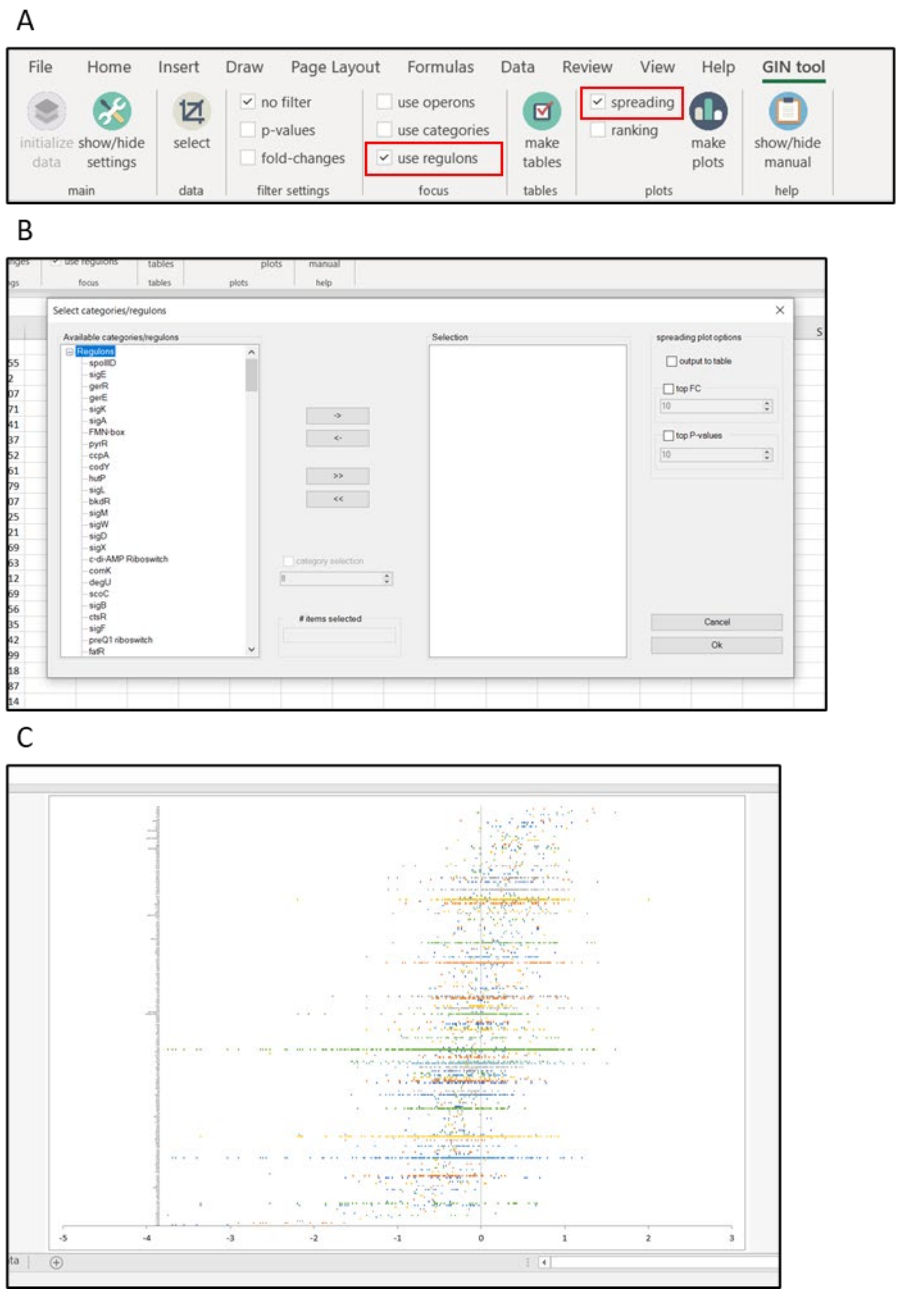


Figure 6: Creating a spread plot of all regulons. A) Selection of focus and type of plot; B) Selection of regulons of interest; C) Spread plot "<u>RegSpreadPlot</u>" with information of fold-changes of all genes in all regulons, in descending order.

Often only the most up- or downregulated regulons are interesting. To show these, go back to "<u>make plots</u>" and tick the "<u>top FC</u>" (fold-change) box and set the number of most strongly regulated regulons to be displayed, *e.g.*, 20 (Figure 7A, red box). By clicking on "<u>Ok</u>" a new sheet is created called "<u>RegSpreadPlotTop20FC</u>", displaying the 20 most affected regulons (Figure 7B). Alternatively, the regulons with the lowest average p-values (see 10) can be selected (Figure 7A, red box).

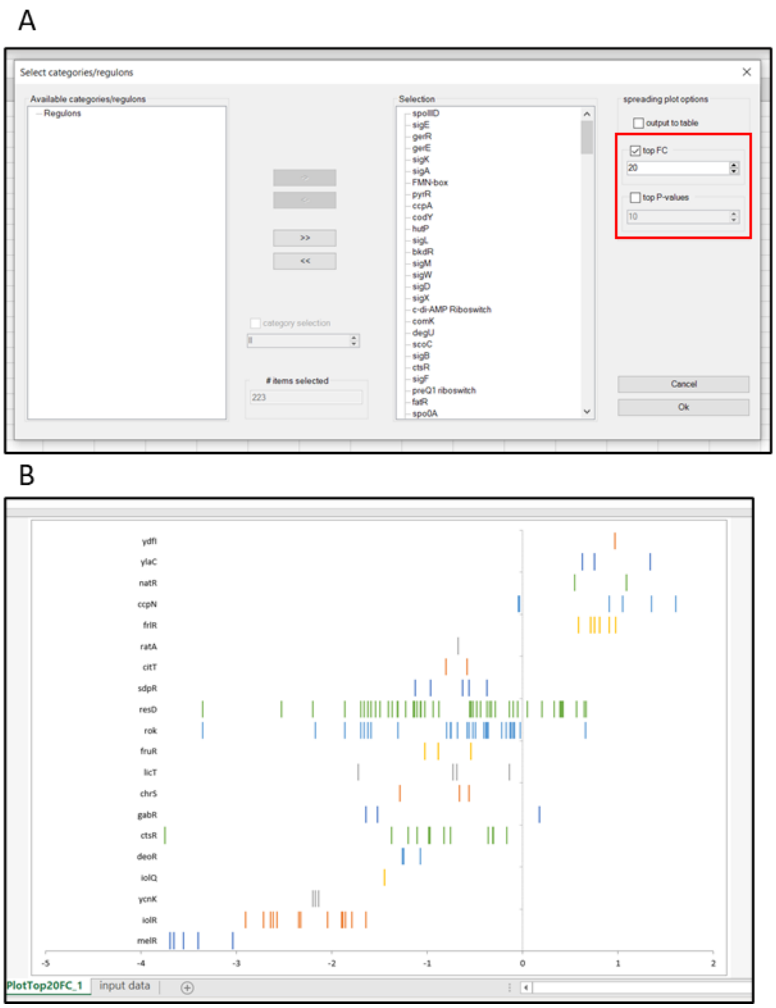


Figure 7: Creating a spread plot of top 20 affected regulons based on average fold-changes. A) Selection of spreading plot options; B) Plot of top 20 most affected regulons based on average FC "RegSpreadPlotTop20FC".

As default the most upregulated regulons are displayed on top. This order can be reversed. For this, click "show/hide settings" and the "sort direction" window becomes visible (Figure 8A, red box). Tick "ascending", click on "show/hide settings" and "make plots", select the regulons of interest, if necessary also select the best scoring regulons using the FC or p-value selection, and click "Ok". Figure 8B shows the reversed result of Figure 6C.

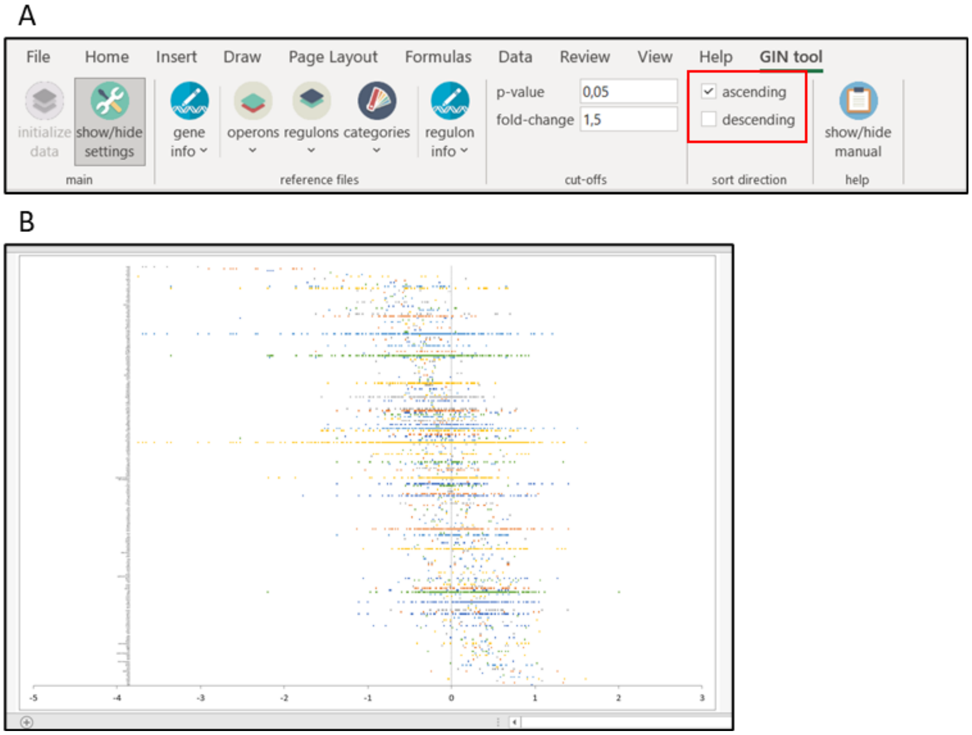


Figure 8: Creating a spread plot of all regulons in ascending order. A) In settings bar, ticking direction in "sort direction"; B) Spread plot "RegSpreadPlot" with information of fold-changes of all genes in all regulons, in ascending order.

It can be useful to know what genes are part of the selected regulons and what their fold-change and p-values are. To obtain this information click again on "make plots" but this time tick the "output to table" box (Figure 9A, red box). In the example the 10 strongest up or down regulated regulons have been selected. After clicking "Ok" a sheet called "RegSpreadTabTop10FC" is created, listing the selected regulons with the genes that are part of the regulon and the related fold-change (FC) (Figure 9B). Besides, a sheet with the corresponding spread plot called "RegSpreadPlotTop10FC" is created.

A

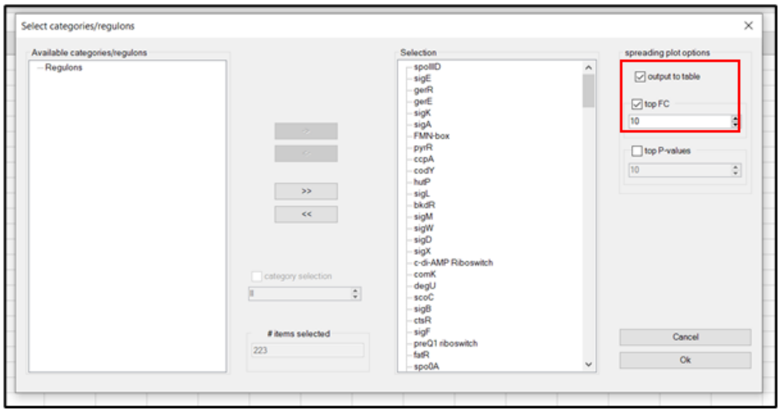


B

| | A | B | C | D |
|---|---|---|---|---|
| 1 | Regulon | Gene | FC | p-value |
| 2 | ydfI | ydfJ | 0.964749 | 0.145385 |
| 3 | ylaC | ylaB | 1.3398 | 0.206763 |
| 4 | ylaC | ylaC | 0.752457 | 0.312897 |
| 5 | ylaC | ylaA | 0.623017 | 0.346254 |
| 6 | chrS | chrS | -0.5587 | 0.223859 |
| 7 | chrS | ywrB | -0.6636 | 0.176841 |
| 8 | chrS | ywrA | -1.28899 | 0.013897 |
| 9 | gabR | gabR | 0.173587 | 0.693414 |
| 10 | gabR | gabD | -1.51829 | 6.2E-05 |
| 11 | gabR | gabT | -1.64061 | 0.000586 |
| 12 | ctsR | ysxC | -0.16648 | 0.635803 |
| 13 | ctsR | lonA | -0.31004 | 0.341778 |
| 14 | ctsR | clpX | -0.31201 | 0.385095 |
| 15 | ctsR | trxA | -0.35974 | 0.321492 |
| 16 | ctsR | disA | -0.7549 | 0.03259 |
| 17 | ctsR | radA | -0.82336 | 0.025296 |

Figure 9B: Generation of table with fold-change and p-value of genes in selected regulons. A) Selection of settings in "Select categories/regulons"; B) Resulting table in sheet "RegSpreadTabTop10FC".

An alternative way to select regulons is by specifying cut-off values for either the fold-change or p-value of the genes that are included in the analysis. Click "show/hide settings" to open the settings bar. The desired cut-offs can be determined in the cut-offs window (Figure 10A, red box). By default, no cut-offs are activated. To use a cut-off, click "show/hide settings" and tick "fold-changes" in the "filter settings" window in the analysis bar (Figure 10B, red box). The plots and tables can be created as described above. For example, the FC cut-off can be changed to 2. After filtering the genes based on a fold-change cut-off of 2, only 14 regulons remain (Figure 10C). The same procedure can be followed to select genes based on p-values, or to select based on both cut-offs, by ticking the appropriate boxes in "filter settings" (Figure 10B).

**!** If the fold-changes in the data set are log2 transformed, then the FC cut-off is also interpreted as such.

**!** The fold-change cut-offs are absolute values, *i.e.*, fold 2 can be both 2-fold downregulation and 2-fold upregulation.

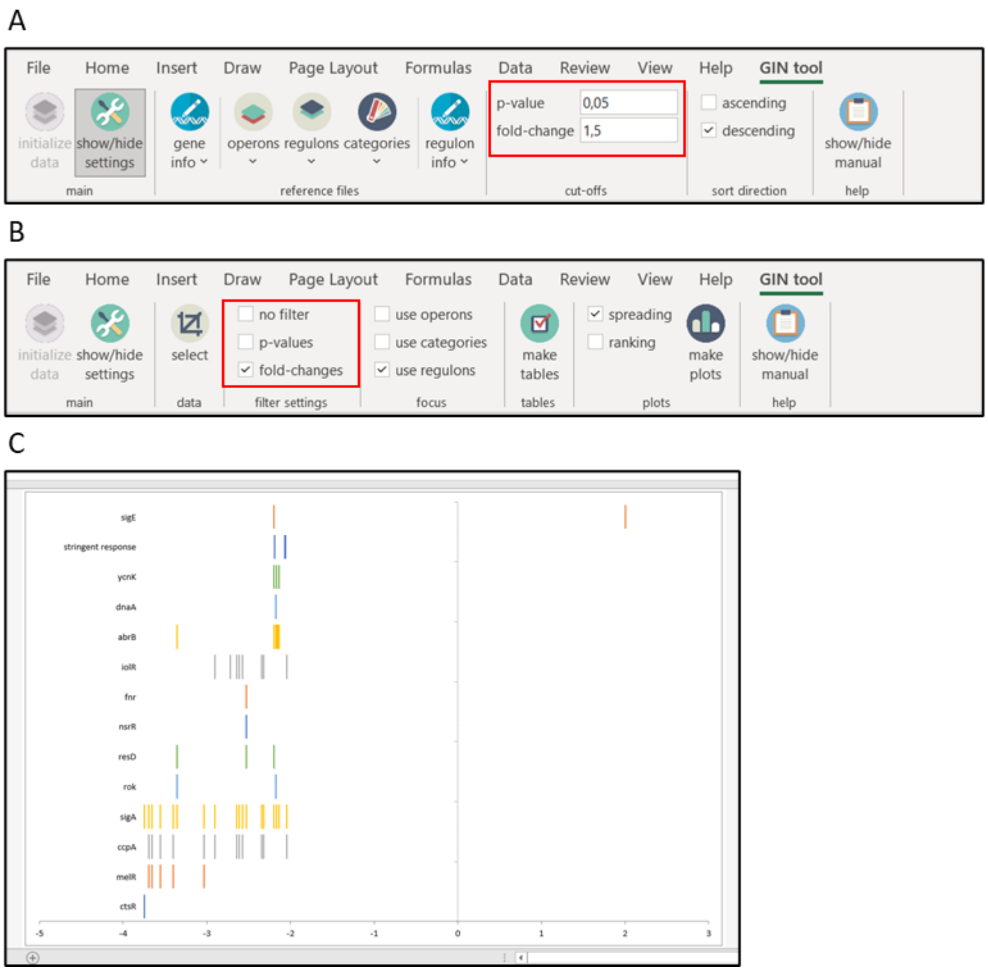


Figure 10: Creating spread plots of regulons after filtering genes. A) Cut-off values can be specified in the "cut-offs" window in the settings bar; B) Cut-offs can be activated by ticking the appropriate boxes in "filter settings" in the analysis bar; C) Example of "RegSpreadPlot" after filtering genes below a fold-change of 2.

#### 6.2 Visual analysis of regulons using bubble plots

The large number of regulons can make it difficult to visually determine how relevant they are. To facilitate this, GINtool can make bubble plots based on the average fold-change of the genes in a regulon (x-axis) and the spreading of fold-changes of genes, defined as the mean absolute deviation (MAD, see 10) of the fold-changes (y-axis). Regulons that show a robust regulation should have fold-changes that are close together, resulting in lower MAD values.

To make these bubble plots, tick "use regulons" in the "focus" window and "ranking" in the "plots" window (Figure 11A, red boxes). Click "make plots" and the "select categories/regulons" window will pop up. Select the relevant regulons as described above (section 6.1). Of note, for this analysis it is not possible to select regulons with e.g. highest fold-change or lowest average p-value, however, fold-change and p-value cut-offs for genes can be activated (Figure 10A). In the example below, no filters are active and all regulons are selected.

After selecting regulons and clicking “Ok”, four sheets are created: “RegRankPlot”, “RegRankPlotBest_v1”, “RegRankPlotBest_v2” and “RegRankTable”. “RegRankPlotBest_v1” and “RegRankPlotBest_v2” will be discussed in section 6.4. An example of “RegRankPlot” is shown in Figure 11B.

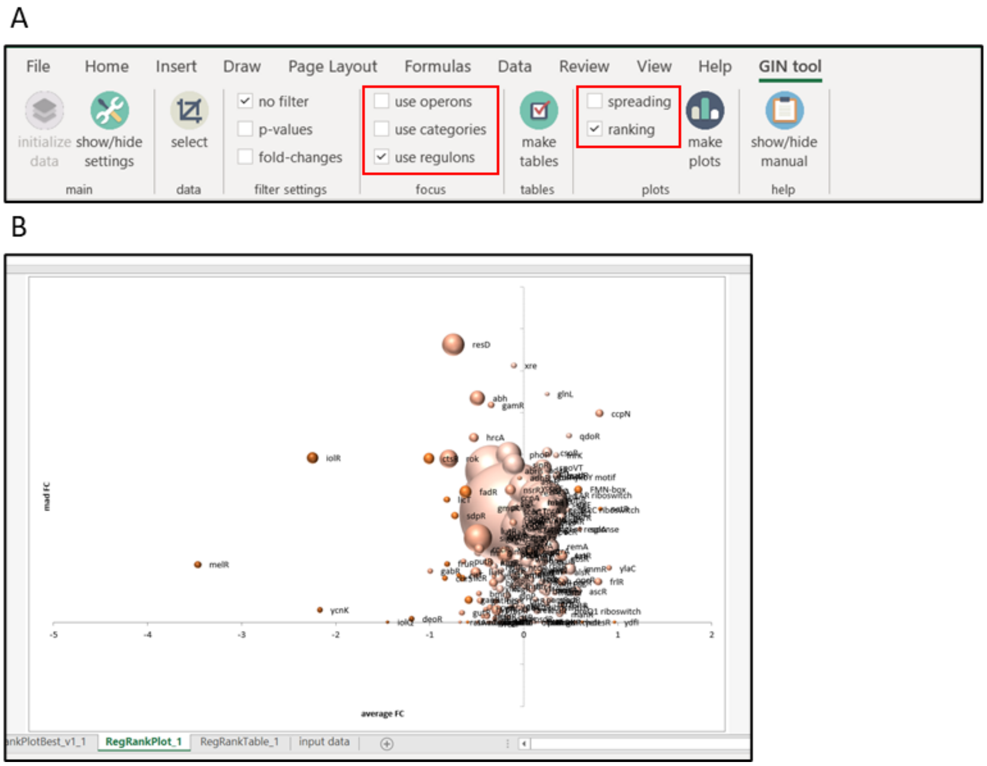


Figure 11: Creating a bubble plot of the average regulation of all regulons. A) Selection of correct settings for creating bubble plots of regulons; B) Example bubble plot “RegRankPlot” of all regulons. Color intensity corresponds to average p-value. Bubble size corresponds to number of genes in regulon.

The use of a bubble plot provides the possibility to also display the size of regulons, *i.e.* the number of genes that are part of a regulon. The example in Figure 11B displays bubbles with different color intensities. The color intensity is related to the average p-value of the genes in the regulon. Regulons with lower average p-values, *i.e.* with more reliable fold-changes, have a more intense color. If bubbles are not required the graph can be easily simplified by chosing another graphical layout in Excel.

The average p-values are distributed in five different classes: average p-value >= 0.5, between 0.5-0.25, between 0.25-0.125, between 0.125-0.0625, and < 0.0625. The classification enables the removal of regulons with high average p-values. Regulons with high p-values can be removed by using the Excel “Chart filter” tool that is activated by clicking on the graph (Figure 12A). In the example in Figure 12A, the regulons with average p-values of >= 0.5 are removed. The bubbles are larger, because Excel works with relative measures for bubble scaling. The scale can be adjusted using the “Format data series” tool that becomes available by right-clicking on a bubble (Figure 12B). In this example, the scale is reduced to 50. Of course, the plots can be converted in other type of graphs and adjusted using standard Excel tools, and the graphs can be exported for furter editing, which can be useful to reposition the names of bubbles or change the size or colors.

A

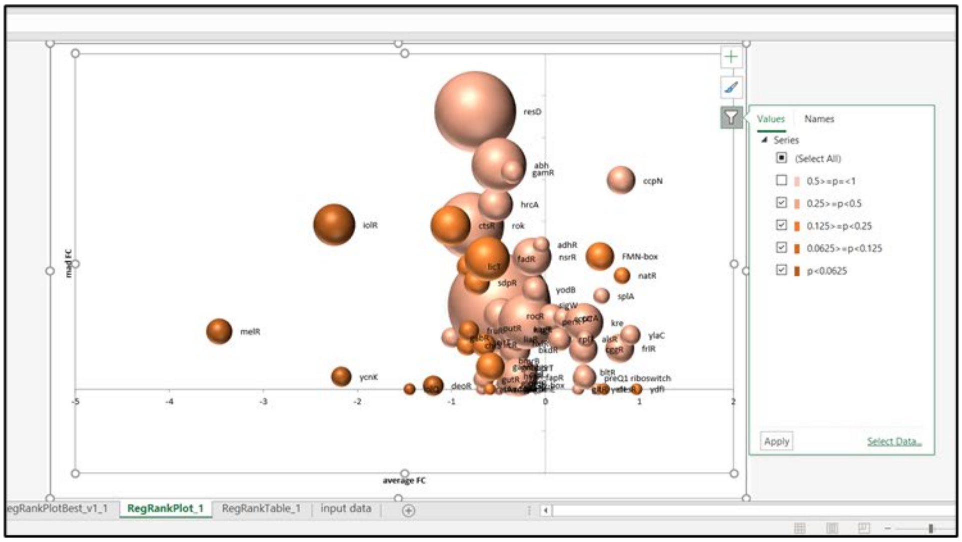

resD
abh
gamR
ccpN
hrcA
iolR
ctsR
rok
adhR
nsrR
FMN-box
fadR
natR
sdpR
yodB
splA
melR
rocR
kre
ylaC
alsR
ycnK
bltR
preQ1 riboswitch
deoR
ydfI
average FC
mad FC
Values
Names
Series
(Select All)
0.5>=p=<1
0.25>=p<0.5
0.125>=p<0.25
0.0625>=p<0.125
p<0.0625
Apply
Select Data...
RegRankPlot_1
RegRankTable_1
input data


B

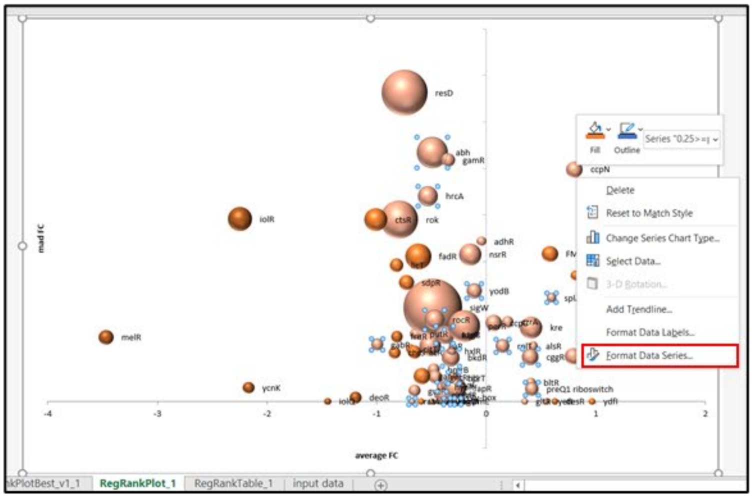

resD
abh
gamR
ccpN
hrcA
iolR
ctsR
rok
adhR
fadR
nsrR
sdpR
yodB
sigW
rocR
kre
melR
alsR
ycnK
bltR
preQ1 riboswitch
deoR
ydfI
average FC
mad FC
Fill
Outline
Delete
Reset to Match Style
Change Series Chart Type...
Select Data...
3-D Rotation...
Add Trendline...
Format Data Labels...
Format Data Series...
RegRankPlot_1
RegRankTable_1
input data


C

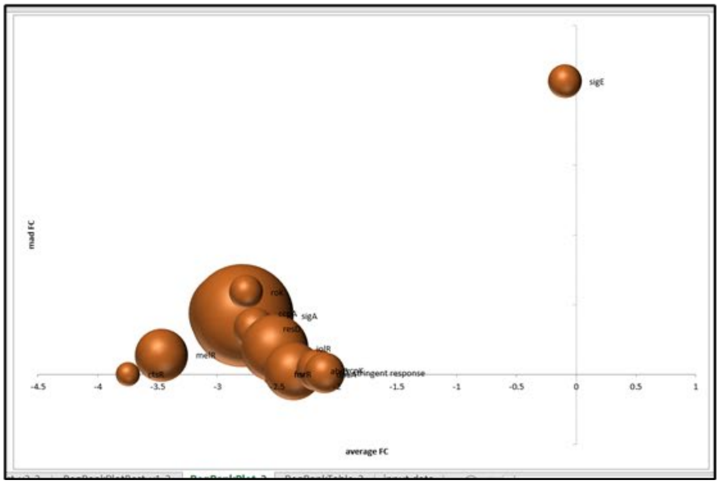

sigE
mad FC
sigA
ctsR
melR
average FC

Figure 12: Creating bubble plots of regulons based on preferences. A) Selecting bubbles based on average p-value of genes in a regulon; B) Rescaling bubble sizes; C) Bubble plot of regulons after filtering genes with FC cut-off of 2.

It is possible display the bubble plots of regulons after filtering the genes in the regulons based on fold-change and p-value (Section 6.1, Figure 10A). When a fold-change cut-off of 2 is active, fewer regulons remain and the bubbles are darker in color, since the average p-values are much lower with this stringent fold-change cut-off value (Figure 12C).

#### 6.3 Regulon evaluation using tables

The average fold-change, MAD- and p-values can also be listed in a table. To do this, click on the "make tables" tab in the "tables" window (Figure 13A, red box). This creates two sheets: "Mapping_details" and "Mapped". "Mapped" will be discussed in section 6.4. The "Mapping_details" sheet contains four different tables (Figure 13B). The first table "Without regulatory directionality" shows the number of genes, average fold-change, MAD fold-change and average p-value for each regulon. The table enables a quick ranking according to any of these parameters, by selecting the headers and using the Excel option "Filter" in the "Sort & Filter" tab in the Home tab (Figure 13C, red box). Of note, when making bubble plots there is also a table created "RegRankTable". This table is the same as the "Mapping_details" table, but then showing only the regulons that were selected.

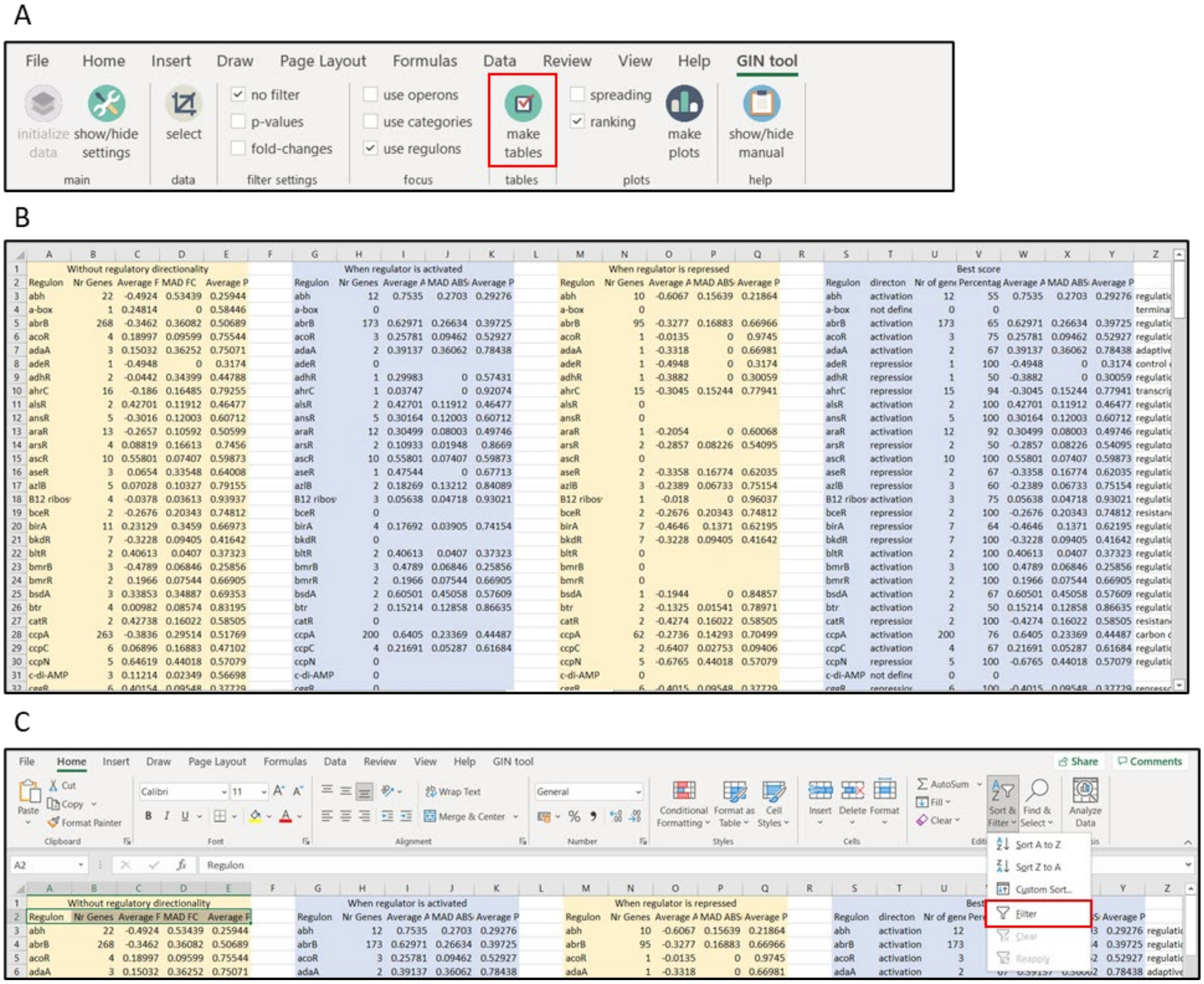


Figure 13: Creating tables with regulon information. A) Selecting the right options for creating tables; B) Sheet "Mapping_details"; C) Adding filters to headers to enable filtering and sorting of the first table.

#### 6.4 Determination of most relevant regulon regulation

The problem with ranking of regulons is that the average fold-change calculation is based on all the genes of a regulon. However, a regulator might repress certain genes and activate others. This bifurcation will reduce the average fold-change. For example, when a regulator significantly activates 5 genes and represses 5 genes then the average fold-change can be close to zero, whereas the regulator/regulon is significantly activated. To correct for this, GINtool uses the regulation direction that has been defined when uploading the regulon file in the "up/down regulation mapping" window discussed in section 4.1 (Figure 4B). This information indicates whether a regulator activates or represses a gene. To incorporate this information, GINtool calculates the situation when the regulator is either activated or repressed, resulting in the second and third table in "Mapping_details" that are labelled "When regulator is activated" and "When regulator is repressed" (Figure 13B). The fourth, most right, table called "Best score", lists the best scoring activity deducted from the second and third tables, based firstly on the number of genes that follow the same type of activity, and in case the numbers are the same, based on the activity with the highest average FC. This final table also indicates the fraction of genes that follow the most relevant regulon activity relative to all the genes that belong to a regulon, indicated in % in column V. This percentage is another important value that indicates how relevant a regulon is for explaining the transcriptome data (see also below).

It is possible that the mode of regulation of a gene has not been defined and therefore could not be selected in the "up/down regulon mapping" window. If this was the case, then the direction of regulon activity in table "Best score" will be 'undefined' (*e.g.*, for regulon 'a-box' in row 4 in Figure 13B). Importantly, when the directionality of gene regulation cannot be defined, then this will also affect the fraction of genes that follow the most relevant regulon activity, and in that case the % in column V can be lower than 50%.

The result of the "Best score" table is automatically displayed as bubble plot when the GINtool option "make plots" is used and the tick box "ranking" is active (see section 6.2). This plot is called "RegRankPlotBest_v1" (Figure 14A). In this plot the negative or positive fold-change indicate whether a regulator is activated or repressed, respectively. *Importantly*(!), this can be a different direction than the measured fold-changes, since an activated repressor (positive FC in the plot) will reduce the expression of genes. That is why in Figure 14A "melR" has a positive value whereas the average fold-change of the MelR regulon was negative as shown in the "RegRankPlot" bubble plot in Figure 11B.

The problem with "RegRankPlotBest_v1" is that it is unclear how many genes of a regulon (columns H and N versus column B in the "Mapping_details" sheet) form the bubble data points. However, the "Best score" table also provides the fraction of the genes that were selected, % in column V, which provides an alternative way to display the most relevant regulons for a quick visual inspection by plotting the average fold-change of the best scoring activity against the fraction (%) of genes in a regulon that are responsible for the calculated fold-change. Such bubble plot is automatically made when the GINtool option "make plots" is used and the tick box "ranking" is on, and is called "RegRankPlotBest_v2" (Figure 14B). In the example, the bubble size has been reduced and regulons with average p-values >= 0.5 are not shown. Since only the most relevant activity of the regulons are chosen for the "Best score" table, the fractions are mostly above 50 %. However, as explained above, the % may be below 50 % when not all genes in a regulon have a defined mode of regulation. Finally, also for this graph type, bubble size and names can be adjusted using standard Excel tools, and the most relevant regulons can also be selected by setting cut-off values in the "filter settings" window as explained in 6.2.

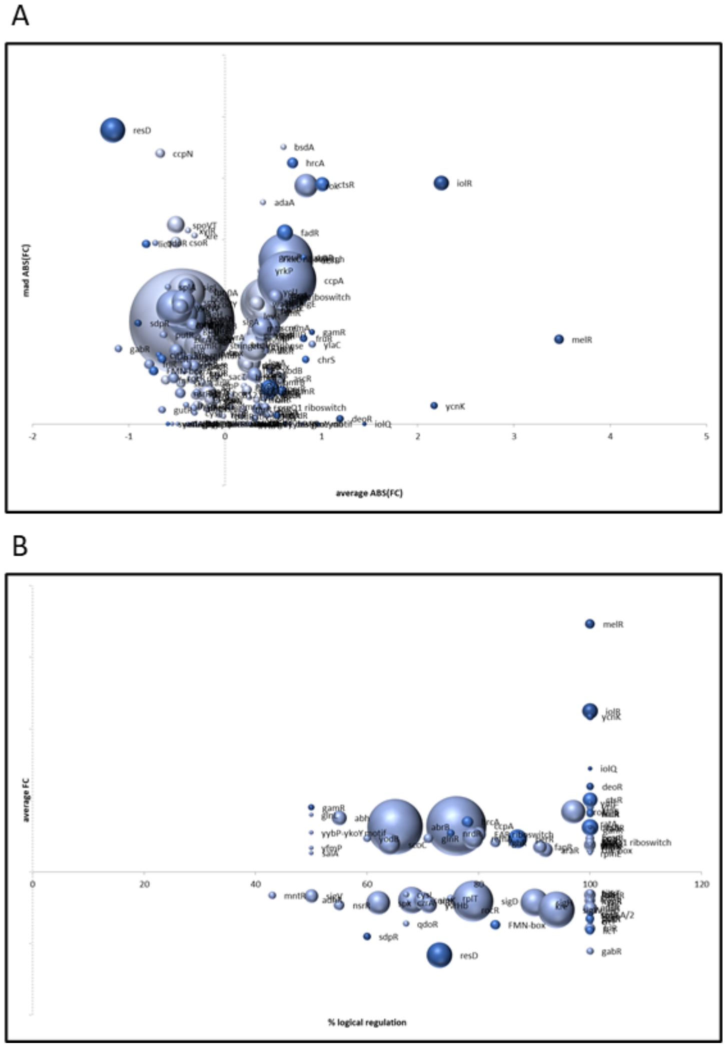


Figure 14: Bubble plots of most relevant regulon activities. A) "RegRankPlotBest_v1" shows the most relevant regulation of each regulon; B) "RegRankPlotBest_v2" shows the fraction of genes that follows the most relevant regulon regulation. Regulons with p-values >= 0.5 are filtered and bubbles are rescaled to 50.

It can be useful to have the "Best score" data of regulons available for every gene. This is possible by using the "make tables" function in GINtool. As described at the beginning of this section, this function produces two table sheets among which one is called "Mapped" (Figure 15).

The table indicates the details of a gene (ID, name, fold-change, p-value, function, description), the number of regulons ("Tot# Regulons"), and names of the regulons to which the gene belongs. In addition, it shows in between brackets the average fold-change (FC) and fraction (%) of genes of the most relevant state of a regulon, based on the "Best score" table in the table sheet "Mapping_details" (e.g., sigE(FC:0.44 71%)). Of note, regulons for which the mode of regulation (activation/repression) could not be indicated will only show the average FC but not the %.

|  | A | B | C | D | E | F | G | H | I |  |
|---|---|---|---|---|---|---|---|---|---|---|
| 1 | Gene id | Gene name | Fold-change | P-value | Function | Description | Tot# Regulons | Regulon_1 | Regulon_2 | Regulo |
| 2 | BSU_11549 | yizD | 2.002516245 | 0.096155044 | unknown | unknown | 1 | sigE(FC:0.44 71%) |  |  |
| 3 | BSU_21030 | yonP | 1.62902704 | 0.227919906 | unknown | unknown | 0 |  |  |  |
| 4 | BSU_14629 | sr1 | 1.601961295 | 0.002606691 | control of arginine m | small regulatory RNA | 2 | sigA(FC:-0.45 60%) | ccpN(FC:-0.83 100%) |  |
| 5 | BSU_36720 | ywmE | 1.504901617 | 0.152470565 | survival of ethanol st | general stress protein | 1 | sigB(FC:-0.39 69%) |  |  |
| 6 | BSU_20300 | yorP | 1.504410801 | 0.181840906 | unknown | unknown | 0 |  |  |  |
| 7 | BSU_06319 | ydzX | 1.503871939 | 0.177236706 | unknown | unknown | 0 |  |  |  |
| 8 | BSU_27809 | yrzT | 1.476434051 | 0.218651856 | unknown | unknown | 0 |  |  |  |
| 9 | BSU_06120 | ydjB | 1.440394618 | 0.039960867 | unknown | unknown | 0 |  |  |  |
| 10 | BSU_20620 | yoqI | 1.409932161 | 0.183178746 | unknown | unknown | 0 |  |  |  |
| 11 | BSU_32090 | yuiA | 1.395148599 | 0.019006795 | unknown | unknown | 2 | codY(FC:-0.38 54%) | tnrA(FC:0.30 57%) |  |
| 12 | BSU_07190 | yezD | 1.392904273 | 0.180524672 | unknown | unknown | 1 | cymR(FC:-0.43 70%) |  |  |
| 13 | BSU_19430 | cdaS | 1.357653606 | 0.145220924 | synthesis of c-di-AMP | sporulation-specific d | 1 | sigG(FC:0.40 64%) |  |  |
| 14 | BSU_29020 | gapB | 1.35051979 | 0.002368569 | anabolic enzyme in gl | glyceraldehyde-3-pho | 2 | ccpN(FC:-0.83 100%) | sigA(FC:-0.45 60%) |  |
| 15 | BSU_14720 | ylaB | 1.339800452 | 0.206763112 | unknown | unknown | 3 | spx(FC:-0.43 62%) | sigA(FC:-0.45 60%) | ylaC(F |
| 16 | BSU_02180 | ybfE | 1.334650537 | 0.051212434 | unknown | unknown | 0 |  |  |  |
| 17 | BSU_26980 | yraE | 1.321525618 | 0.257068695 | unknown | forespore-specific sp | 1 | sigG(FC:0.40 64%) |  |  |
| 18 | BSU_16799 | ylzJ | 1.265454223 | 0.236555726 | control of TepA activi | germination protein, | 2 | spoVT(FC:-0.51 60%) | sigG(FC:0.40 64%) |  |
| 19 | BSU_23270 | ribE | 1.264654964 | 0.002935057 | riboflavin biosynthesi | riboflavin synthase (a | 2 | sigA(FC:-0.45 60%) | FMN-box(FC:-0.74 83%) |  |
| 20 | BSU_36680 | ywmF | 1.231746679 | 0.169942124 | unknown | unknown | 1 | sigB(FC:-0.39 69%) |  |  |
| 21 | BSU_misc_RNA_59 | BSU_misc_RNA_59 | 1.211348333 | 0.192298838 |  |  | 0 |  |  |  |
| 22 | BSU_28260 | leuC | 1.20879374 | 0.000917827 | biosynthesis of leucin | 3-isopropylmalate de | 4 | sigA(FC:-0.45 60%) | tnrA(FC:0.30 57%) | codY(F |
| 23 | BSU_22849 | ypzH | 1.157034029 | 0.301486735 | unknown | unknown | 0 |  |  |  |
| 24 | BSU_11290 | yjaV | 1.115583996 | 0.241513723 | [[category\|Sporulatic | [[protein\|SigE]]-depe | 1 | sigE(FC:0.44 71%) | | |
| 25 | BSU_18899 | yozX | 1.099514197 | 0.317003651 | unknown | unknown | 0 |  |  |  |
| 26 | BSU_26380 | yqaB | 1.098258018 | 0.214665205 | unknown | similar to phage-relat | 0 |  |  |  |

Figure 15: Overview of the table in sheet named "Mapped". Regulon information is shown per gene. Regulon FC 'NaN' indicates the regulon directionality is unknown.

#### 7. Analysis of functional categories

*First follow the steps described in **section 4**.*

7.1 Visual analysis of functional categories using spread plots

GINtool gives the option to rank functional categories based on the average fold-change of all genes in a functional category. The distribution of fold-changes of genes of functional categories can be visualized in a spread plot. Start by ticking "use categories" as "focus" and "spreading" for "plots" (Figure 16A). Click "make plots", the "Select categories/regulons" window will pop up (Figure 16B).

By clicking on "Categories" in the left window "Available categories/regulons", the subcategories become visible. Categories and subcategories can be selected and moved to the right window "Selection" by clicking on the single arrow button. It is also possible to select different levels of categories by ticking the "category selection" box, indicating the level you would like to display and subsequently clicking the double arrow button (Figure 16B, red box). The most detailed categories, are levels III and IV for the *B. subtilis* annotation. To produce the spread plot, click "Ok". This will create a new sheet called "CatSpreadPlot", displaying the fold-changes of all genes in the selected categories, whereby the categories are ranked based on the average fold-change (Figure 6C). The x-axis indicates the fold-change.

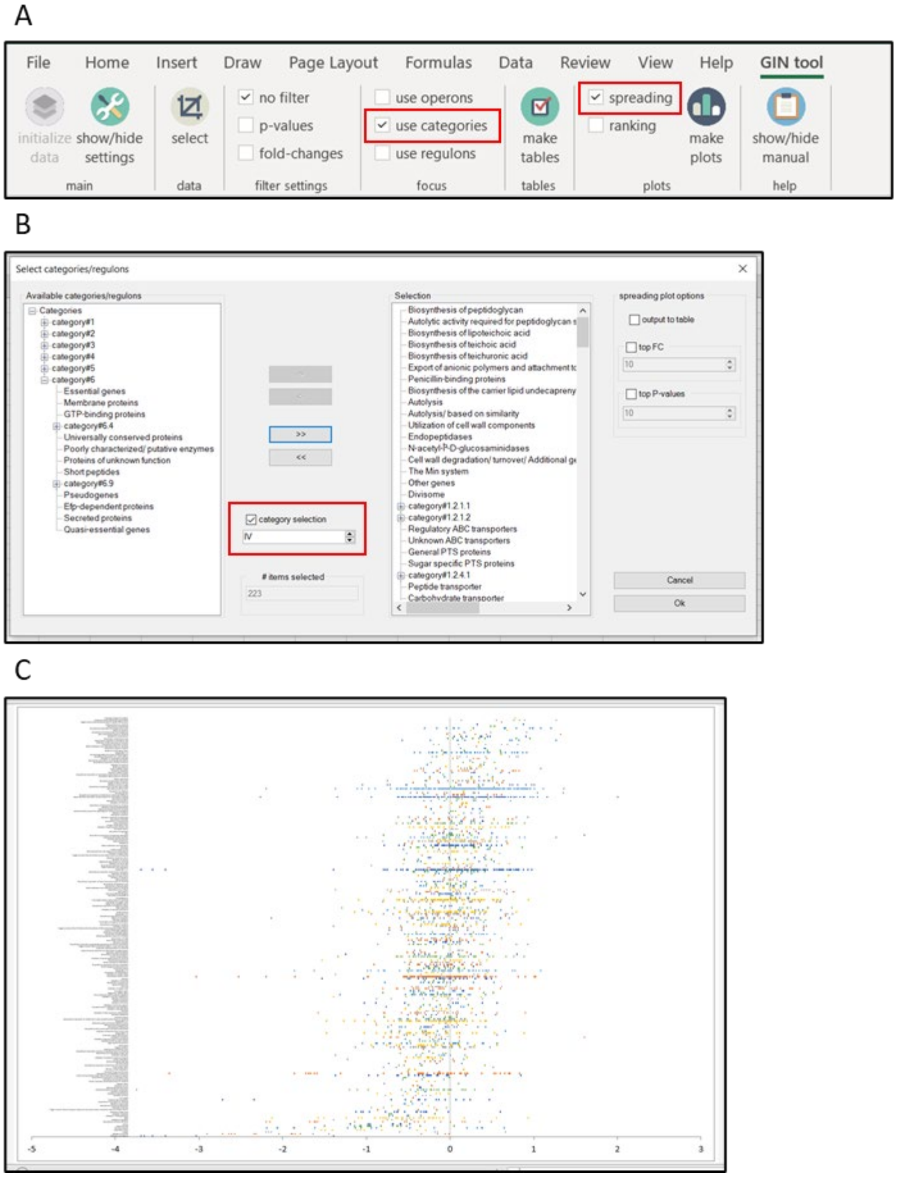


Figure 16: Creating a spread plot of functional categories. A) Selection of focus and type of plot; B) Selection of categories of interest; C) Spread plot "CatSpreadPlot" with information of fold-changes of all genes in the selected categories, in descending order.

Often only the most up- or down-regulated categories are interesting. To show these, go back to "make plots" and tick the "top FC" (fold-change) box and set the number of most strongly regulated categories to be displayed, *e.g.*, 20 (Figure 17A, red box). By clicking on "Ok" a new sheet is created called "CatSpreadPlotTop20FC", displaying the 20 most affected selected categories (Figure 17B). Alternatively, the categories with the lowest average P-values can be selected (Figure 17A, red box).

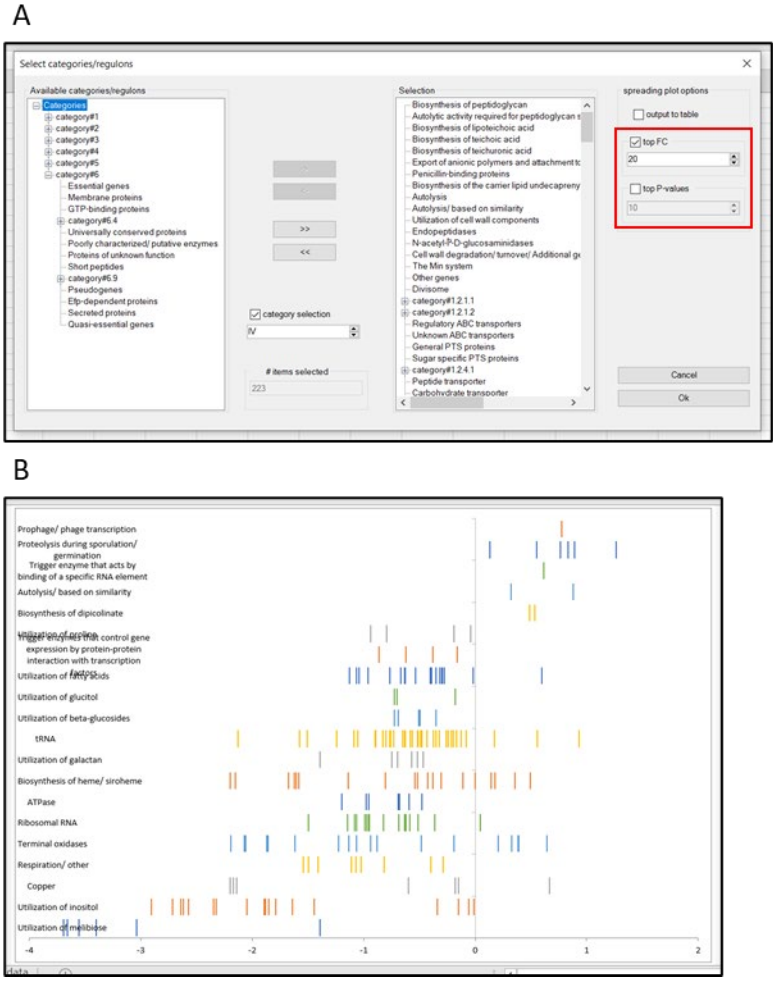


Figure 17: Creating a spread plot of top 20 affected categories based on average fold-changes. A) Selection of spreading plot options; B) Plot of top 20 most affected selected categories based on average FC "CatSpreadPlotTop20FC".

As default the most upregulated categories are displayed on top. This order can be reversed. For this, click on the "show/hide settings" tab so that the "sort direction" window becomes visible (Figure 18A, red box). Tick "ascending", click on "show/hide settings" and "make plots", select the categories of interest, if necessary also select the best scoring categories using the FC or p-value selection, and click on "Ok". Figure 18B shows the reversed result of Figure 16C.

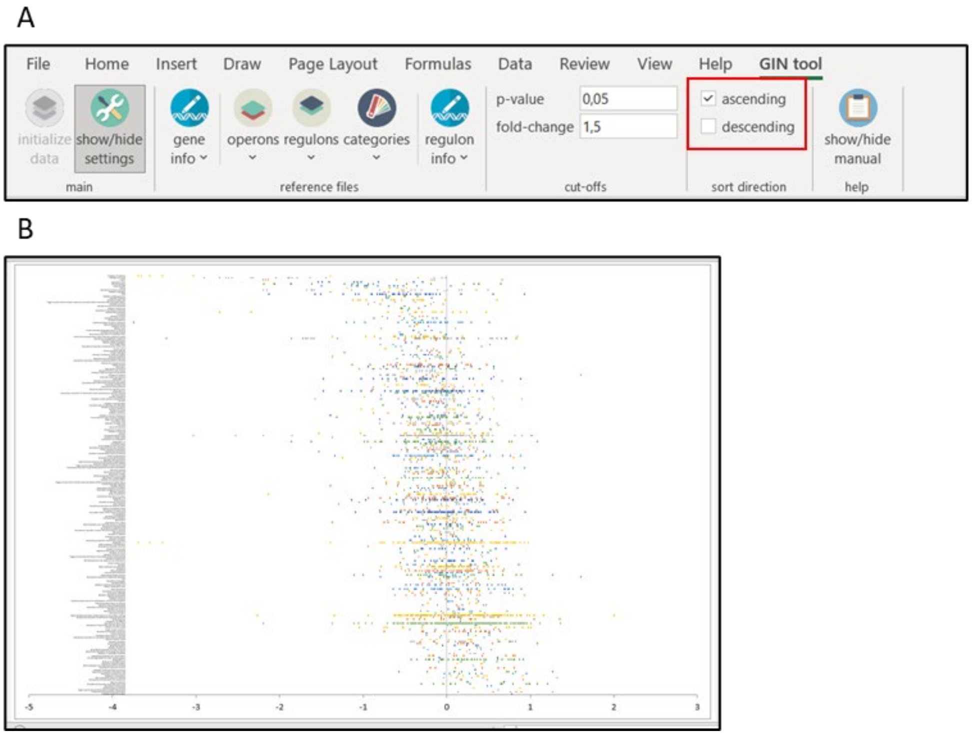


Figure 18: Creating a spread plot of all selected categories in ascending order. A) In settings bar, tick direction in "sort direction"; B) Spread plot "CatSpreadPlot" with information of fold-changes of all genes in the selected categories, in ascending order.

It can be useful to know what genes are part of the selected categories and what their fold-change and p-values are. To obtain this information click again on "make plots" but this time tick the "output to table" box (Figure 19A, red box). In the example the 10 strongest up- or down regulated categories have been selected. After clicking "Ok" a sheet called "CatSpreadTabTop10FC" is created, listing the selected categories with the genes that are part of the category and the related fold-change (FC) (Figure 19B). Besides, a sheet with the corresponding spread plot called "CatSpreadPlotTop10FC" is created.

A

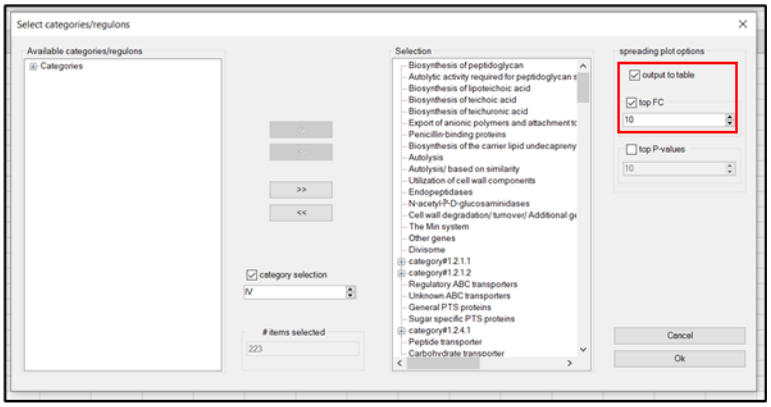


B

| | A | B | C | D |
|---|---|---|---|---|
| 1 | Category | Gene | FC | p-value |
| 2 | Copper | ycnJ | -2.13901 | 4.12E-05 |
| 3 | Copper | ycnI | -2.16938 | 2.27E-05 |
| 4 | Copper | ycnK | -2.1989 | 8.18E-05 |
| 5 | Biosynthesis of heme/ sir | ctaO | -2.15349 | 0.000192 |
| 6 | Biosynthesis of heme/ sir | ctaA | -2.19659 | 3.75E-06 |
| 7 | Sporulation proteins/ oth | ctaA | -2.19659 | 3.75E-06 |
| 8 | Utilization of inositol | iolG | -2.04504 | 3.79E-05 |
| 9 | Utilization of inositol | iolE | -2.32272 | 1.43E-06 |
| 10 | Utilization of inositol | iolF | -2.34329 | 3.1E-06 |
| 11 | Utilization of inositol | iolD | -2.57104 | 9.31E-08 |
| 12 | Utilization of inositol | iolB | -2.61435 | 1.13E-07 |
| 13 | Utilization of inositol | iolC | -2.64146 | 4.25E-08 |
| 14 | Utilization of inositol | iolT | -2.71456 | 0.000894 |
| 15 | Utilization of inositol | iolA | -2.9014 | 4.57E-09 |
| 16 | Carbohydrate transporte | iolF | -2.34329 | 3.1E-06 |
| 17 | Carbohydrate transporte | iolT | -2.71456 | 0.000894 |

Figure 19B: Generation of table with fold-change and p-value of genes in selected categories. A) Selection of settings in "Select categories/regulons"; B) Resulting table in sheet "CatSpreadTabTop10FC".

An alternative way to select categories is by specifying cut-off values for either the fold-change or p-value of the genes that are included in the analysis. Click "show/hide settings" to open the settings bar. The desired cut-offs can be determined in the cut-offs window (Figure 20A, red box). By default, no cut-offs are activated. To use a cut-off, click "show/hide settings" and tick "fold-changes" in the "filter settings" window in the analysis bar (Figure 20B, red box). The plots and tables can be created as described above. For example, the FC cut-off can be changed to 2. After filtering the genes based on a fold-change cut-off of 2, only 14 categories remain (Figure 20C). The same procedure can be followed to select genes based on p-values, or to select based on both cut-offs, by ticking the appropriate boxes in "filter settings" (Figure 20B).

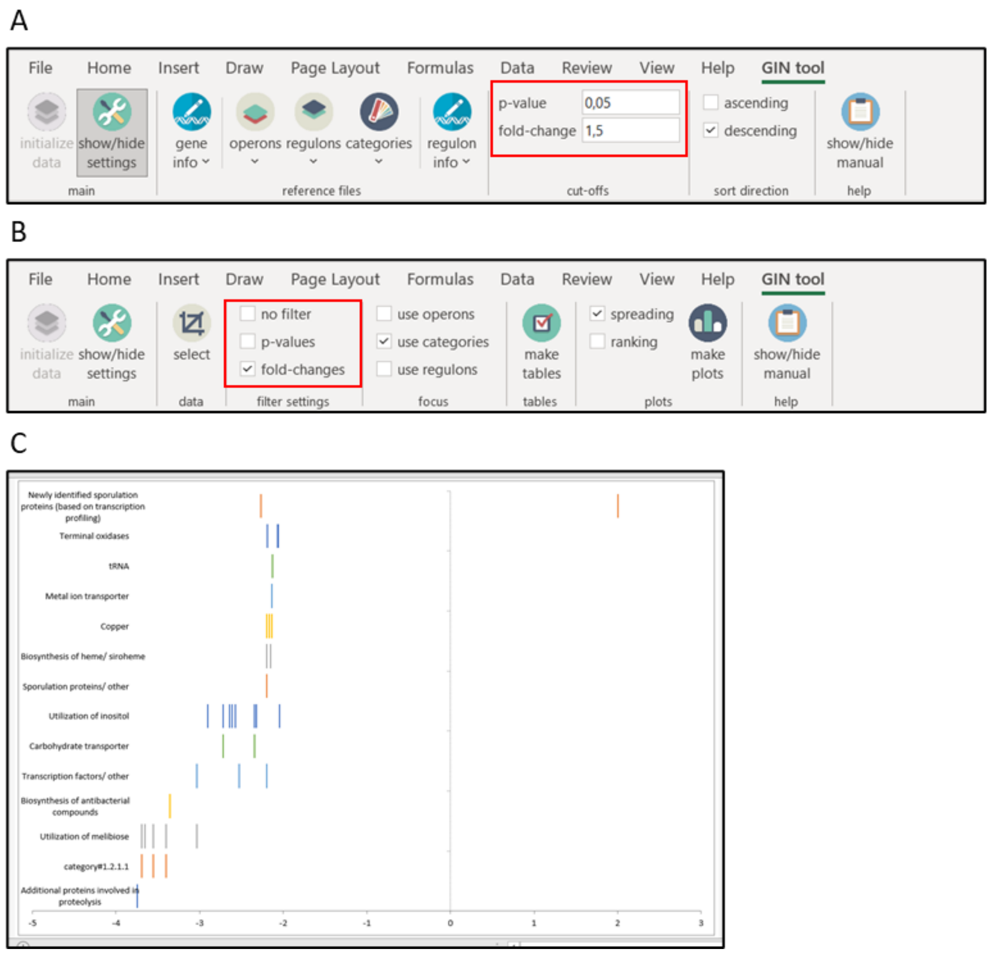


Figure 20: Creating a spread plots of categories after filtering genes. A) Cut-off values can be specified in the "cut-offs" window in the settings bar; B) Cut-offs can be activated by ticking the appropriate boxes in "filter settings" in the analysis bar; C) Example of "CatSpreadPlot" after filtering genes below a fold-change of 2.

**!** If the fold-changes in the data set are log2 transformed, then the FC cut-off is also interpreted as such.

**!** The fold-change cut-offs are absolute values, *i.e.*, fold 2 can be both 2-fold downregulation and 2-fold upregulation.

#### 7.2 Visual analysis of functional categories using bubble plots

The large number of categories can make it difficult to visually determine how relevant they are. To facilitate this, GINtool can make bubble plots based on the average fold-change of the genes in a category (x-axis) and the spreading of fold-changes of genes, defined as the mean absolute deviation (MAD) of the fold-changes (y-axis). Categories that show a robust regulation should have fold-changes that are close together, resulting in lower MAD values.

To make these bubble plots, tick "use categories" in the "focus" window and "ranking" in the "plots" window (Figure 21A, red boxes). Click "make plots" and the "select categories/regulons" window will pop up. Select the relevant categories as described above (section 7.1). Of note, for this analysis it is not possible to select categories with the highest fold-change or lowest average p-value, but filters for genes can be activated (Figure 20A). In this example, no filters are active and the IV$^{th}$ category level is selected.

After selecting categories and clicking "Ok", four sheets are created: "CatRankPlot", "CatRankPlotBest_v1", "CatRankPlotBest_v2" and "CatRankTable". "CatRankTable" is the same table as "Mapping details" which will be discussed in section 7.3, "CatRankPlotBest_v1" and "CatRankPlotBest_v2" will be discussed in section 7.4. An example of "CatRankPlot" is shown in Figure 21B.

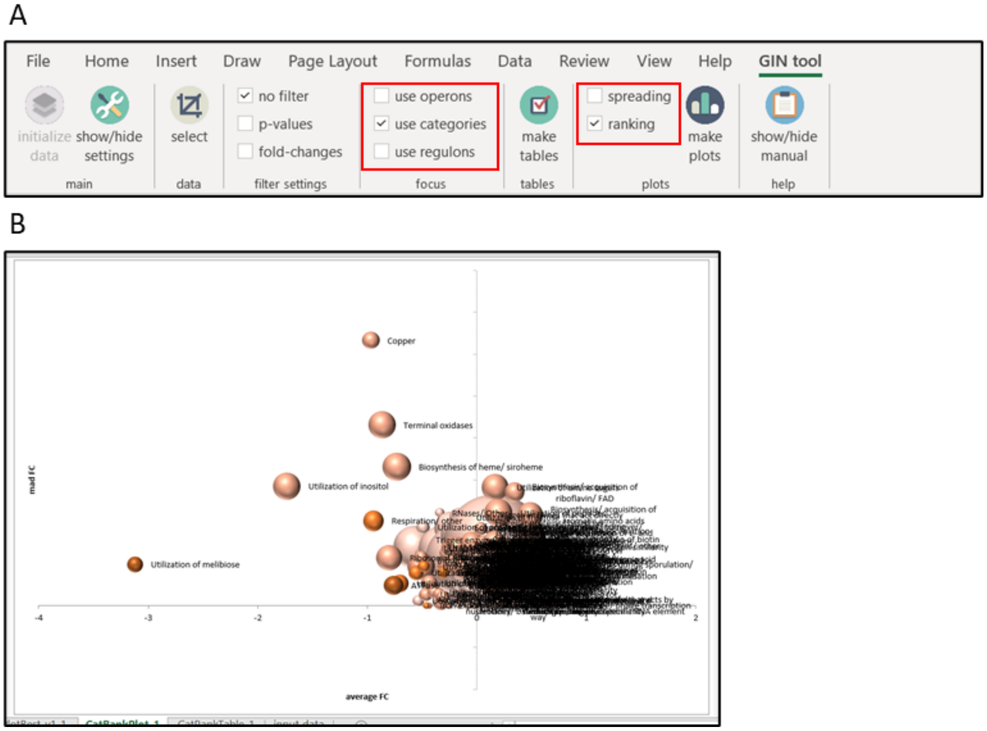


Figure 21: Creating a bubble plot of the average regulation of all selected categories. A) Selection of correct settings for creating bubble plots of categories; B) Example bubble plot "CatRankPlot" of all included categories and genes. Color intensity corresponds to average p-value. Bubble size corresponds to number of genes in regulon.

The use of a bubble plot provides the possibility to also display the size of categories, *i.e.*, the number of genes of a category. The example in Figure 21B displays bubbles with different color intensities. The color intensity is related to the average p-value of the genes in the category. Categories with lower average p-values, *i.e.*, with more reliable fold-changes, have a more intense color.

The average p-values are distributed in five different classes: average p-value >= 0.5, between 0.5-0.25, between 0.25-0.125, between 0.125-0.0625, and < 0.0625. The classification enables the removal of categories with high average p-values. Categories with high p-values can be removed by using the Excel "Chart filter" tool that is activated by clicking on the graph (Figure 22A). In the example in Figure 22A, the categories with average p-values of >= 0.5 are removed. The bubbles are larger, because Excel works with relative measures for bubble scaling. The scale can be adjusted using the "Format data series" tool that becomes available by right-clicking on a bubble (Figure 22B). In this example, the scale is reduced to 50. Of course, the plots can be converted and adjusted using other standard Excel tools, and the graphs can be exported for furter editing, which can be useful to reposition the names of bubbles or change the colors.

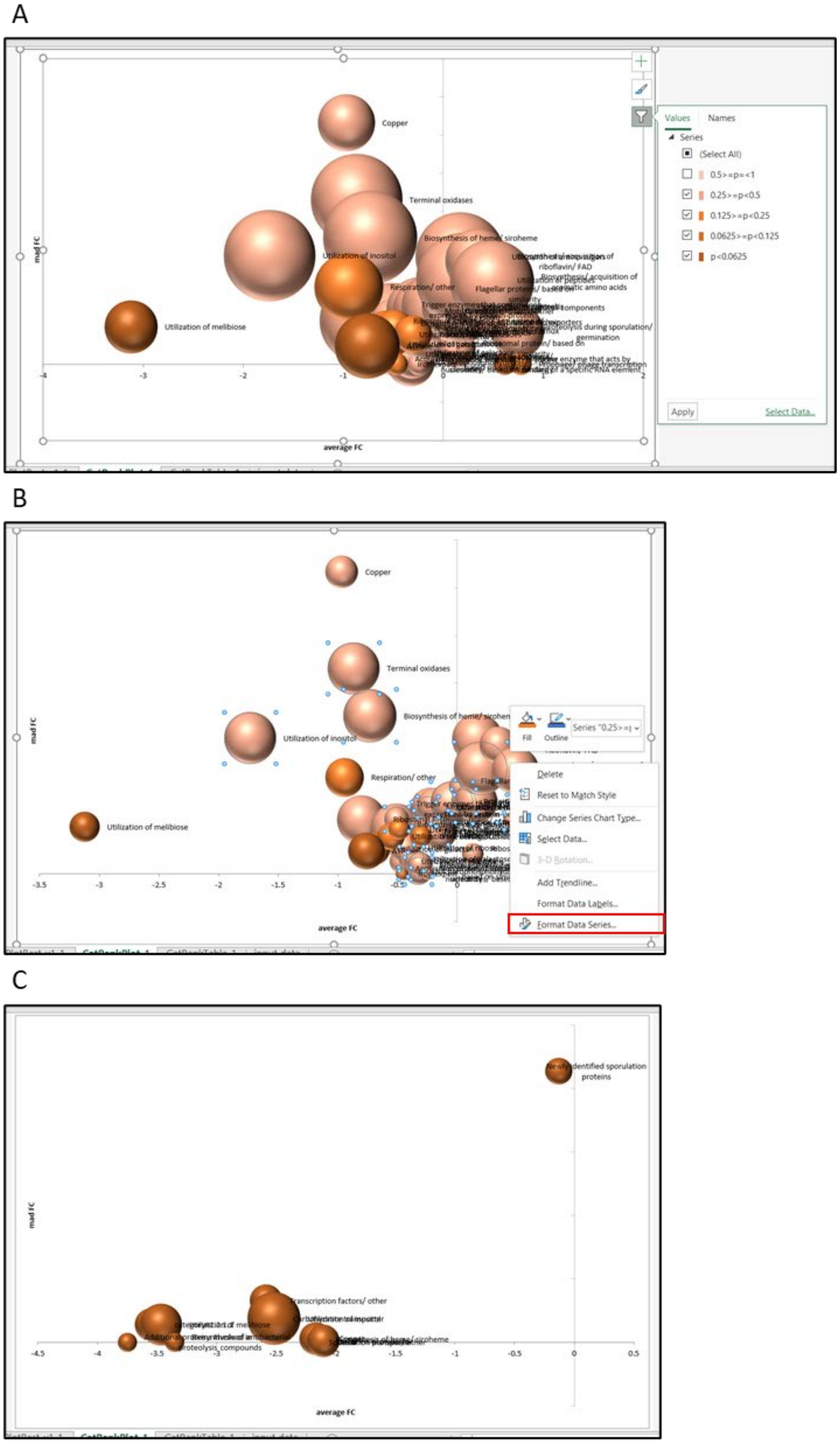


Figure 22: Creating bubble plots of categories based on preferences. A) Selecting bubbles based on average p-value of genes in a category; B) Rescaling bubble sizes; C) Bubble plot of categories after filtering genes with FC cut-off of 2.

It is possible to display the bubble plots of categories after filtering the genes in the categories based on fold-change and p-value (Section 7.1, Figure 20A). When a fold-change cut-off of 2 is active, fewer categories remain and the bubbles are darker in color, since the average p-values are much lower with this stringent fold-change cut-off value (Figure 22C).

#### 7.3 Functional category evaluation using tables

The average fold-change, MAD- and p-values can also be listed in a table. To do this, click on the "make tables" tab in the "tables" window (Figure 23A, red box). This creates two sheets: "Mapping_details" and "Mapped". "Mapped" will be discussed in section 7.4. The "Mapping_details" sheet contains four different tables (Figure 23B). The first table "Plot data" shows the number of genes, average fold-change, MAD fold-change and average p-value for each regulon. The table enables a quick ranking according to any of these parameters, by selecting the headers and using the Excel option "Filter" in the "Sort & Filter" tab in the Home tab (Figure 23C, red box). Of note, when making bubble plots there is also a table created "CatRankTable". This table is the same as the "Mapping_details" table, but then showing only the categories that were selected.

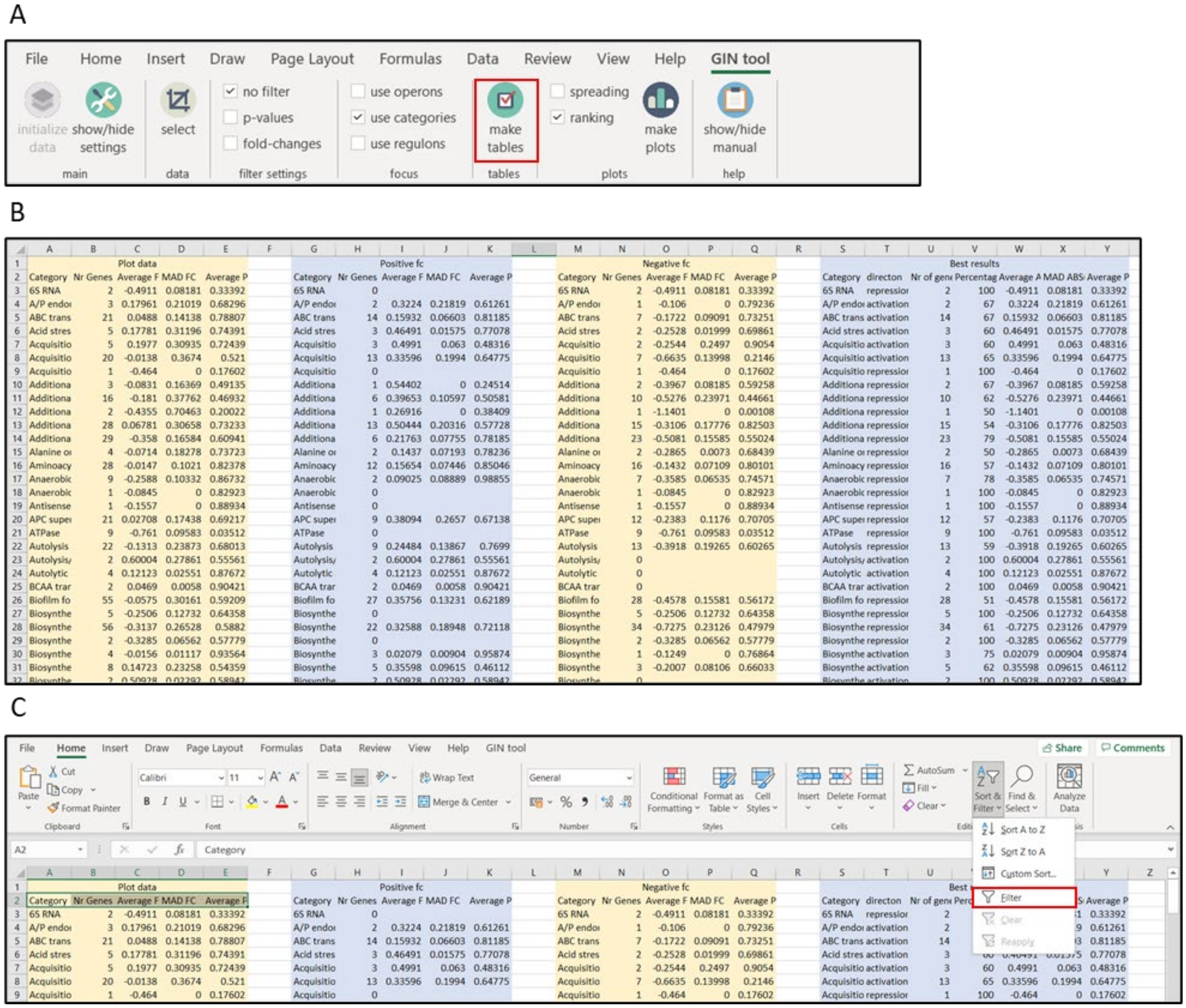


Figure 23: Creating tables with functional category information. A) Selecting the right options for creating tables; B) Sheet "Mapping_details"; C) Adding filters to headers to enable filtering and sorting of the first table.

#### 7.4 Determination of most relevant functional category regulation

The problem with ranking of categories is that the average fold-change calculation is based on all the genes of a category. However, it is possible that some genes are activated whereas others are repressed. If this heterogeneity is the case, then the related category is less relevant. To identify categories that show the most consistent activation or repression, GINtool can calculate the number of genes of a category that corresponds to either activation or repression of the category. This is automatically performed when "Mapping_details" is created (see section 7.3). This is reflected in the second and third table in "Mapping_details" that are labelled "Positive Fc" and "Negative Fc". The fourth, most right table called "Best results", lists the best scoring activity of the categories from the second and third tables, based firstly on the number of genes that follow the same type of activity, and in case the numbers are the same, based on the activity with the highest average FC. This final table also indicates the fraction of genes that follow the most relevant category activity relative to all the genes that belong to the category, indicated in % in column V.

The result of the "Best results" table is automatically displayed as bubble plot when the GINtool option "make plots" is used and the tick box "ranking" is active (see section 7.2). This plot is called "CatRankPlotBest_v1" (Figure 24A).

The problem with "CatRankPlotBest_v1" is that it is unclear how many genes of a category (columns H and N versus column B in the "Mapping_details" sheet) form the bubble data points. However, the fourth "Best results" table also provides the fraction of the genes that were selected, % in column V, which provides an alternative way to display the most relevant categories for a quick visual inspection by plotting the average fold-change of the best scoring activity against the fraction (%) of genes in a category that are responsible for the calculated fold-change. Such bubble plot is automatically made when the GINtool option "make plots" is used and the tick box "ranking" is on, and is called "CatRankPlotBest_v2" (Figure 24B). In the example, the bubble size has been reduced and categories with average p-values >= 0.5 are not shown. Since only the most relevant activity of the categories are chosen, the fraction is always above 50 %. Finally, also for this graph type, bubble size and names can be adjusted using standard Excel tools, and the most relevant categories can also be selected by setting cut-off values in the "filter_settings" window as explained in 7.2.

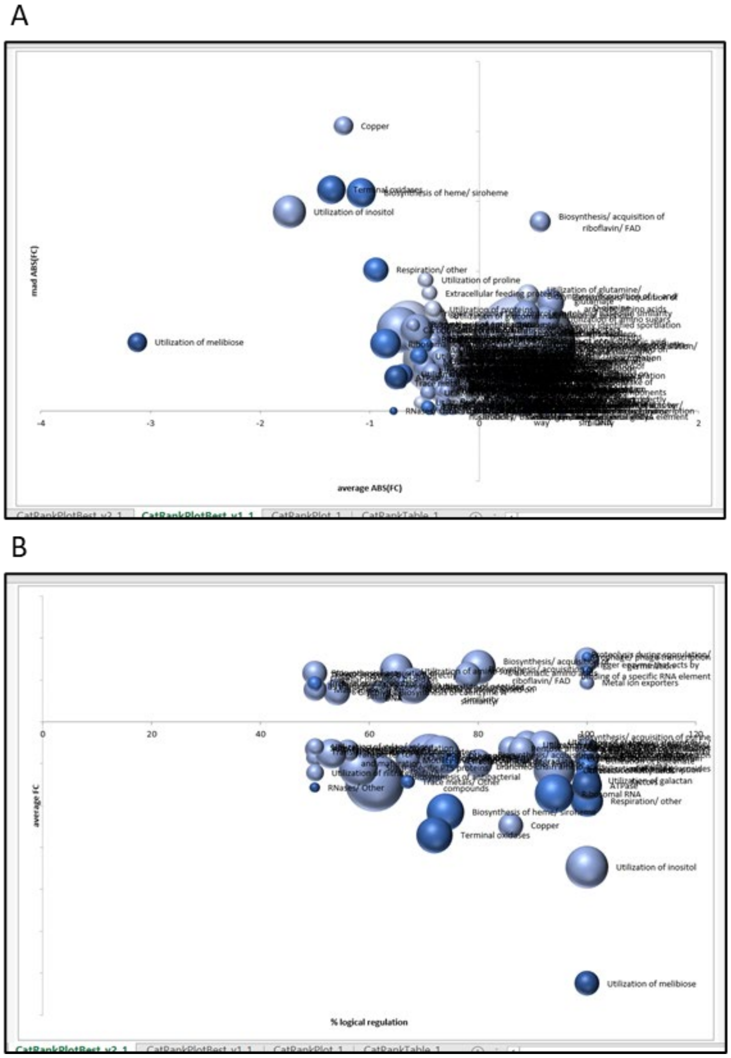


Figure 24: Bubble plots of most relevant category activities. A) "CatRankPlotBest_v1" shows the most relevant regulation of each selected category; B) "CatRankPlotBest_v2" shows the fraction of genes that follows the most relevant category regulation. Categories with p-values >= 0.5 are filtered and bubbles are rescaled to 50.

It can be useful to have the "Best results" data of categories available for every gene. This is possible by using the "make tables" function in GINtool. As described at the beginning of this section, this function produces two table sheets among which one called "Mapped" (Figure 25).

The table indicates the details of a gene (ID, name, fold-change, p-value, function, description), the number of categories ("Tot# Categories"), and names of the categories to which the gene belongs. In addition, it shows in between brackets the average fold-change (FC) and fraction (%) of genes of the most relevant state of a category based on the "Best results" table in the table sheet "Mapping_details" (e.g., SP-beta prophage(FC:0.48 54%)).

| | A | B | C | D | E | F | G | H | I | |
|---|---|---|---|---|---|---|---|---|---|---|
| 1 | Gene id | Gene name | Fold-change | P-value | Function | Description | Tot# Categories | Category_1 | Category_2 | Cat |
| 2 | BSU_11549 | yizD | 2.002516245 | 0.096155044 | unknown | unknown | 2 | Newly identified spor | Membrane proteins(FC:-0 | |
| 3 | BSU_21030 | yonP | 1.62902704 | 0.227919906 | unknown | unknown | 1 | SP-beta prophage(FC:0.48 54%) | | |
| 4 | BSU_14629 | sr1 | 1.601961295 | 0.002606691 | control of arginine m | small regulatory RNA | 3 | Biosynthesis/ acquisit | Utilization of arginine | Reg |
| 5 | BSU_36720 | ywmE | 1.504901617 | 0.152470565 | survival of ethanol st | general stress protein | 2 | General stress protei | Membrane proteins(FC:-0 | |
| 6 | BSU_20300 | yorP | 1.504410801 | 0.181840906 | unknown | unknown | 1 | SP-beta prophage(FC:0.48 54%) | | |
| 7 | BSU_06319 | ydzX | 1.503871939 | 0.177236706 | unknown | unknown | 1 | Proteins of unknown function(FC:-0.35 60%) | | |
| 8 | BSU_27809 | yrzT | 1.476434051 | 0.218651856 | unknown | unknown | 1 | Proteins of unknown function(FC:-0.35 60%) | | |
| 9 | BSU_06120 | ydjB | 1.440394618 | 0.039960867 | unknown | unknown | 2 | Prophage 3(FC:-0.30 ( | Proteins of unknown func | |
| 10 | BSU_20620 | yoql | 1.409932161 | 0.183178746 | unknown | unknown | 1 | SP-beta prophage(FC:0.48 54%) | | |
| 11 | BSU_32090 | yuiA | 1.395148599 | 0.019006795 | unknown | unknown | 1 | Proteins of unknown function(FC:-0.35 60%) | | |
| 12 | BSU_07190 | yezD | 1.392904273 | 0.180524672 | unknown | unknown | 1 | Proteins of unknown function(FC:-0.35 60%) | | |
| 13 | BSU_19430 | cdaS | 1.357653606 | 0.145220924 | synthesis of c-di-AMP | sporulation-specific d | 2 | Metabolism of signall | Sporulation proteins/ oth | |
| 14 | BSU_29020 | gapB | 1.35051979 | 0.002368569 | anabolic enzyme in gl | glyceraldehyde-3-pho | 1 | Gluconeogenesis(FC:0.58 50%) | | |
| 15 | BSU_14720 | ylaB | 1.339800452 | 0.206763112 | unknown | unknown | 2 | Membrane proteins(F | Proteins of unknown func | |
| 16 | BSU_02180 | ybfE | 1.334650537 | 0.051212434 | unknown | unknown | 2 | Membrane proteins(F | Proteins of unknown func | |
| 17 | BSU_26980 | yraE | 1.321525618 | 0.257068695 | unknown | forespore-specific sp | 1 | Spore coat protein/ based on similarity(FC:0.54 | | |
| 18 | BSU_16799 | ylzJ | 1.265454223 | 0.236555726 | control of TepA activi | germination protein, | 2 | Proteolysis during spc | Additional germination pr | |
| 19 | BSU_23270 | ribE | 1.264654964 | 0.002935057 | riboflavin biosynthesi | riboflavin synthase (a | 1 | Biosynthesis/ acquisition of riboflavin/ FAD(FC:( | | |
| 20 | BSU_36680 | ywmF | 1.231746679 | 0.169942124 | unknown | unknown | 2 | General stress protei | Membrane proteins(FC:-0 | |
| 21 | BSU_misc_RNA_59 | BSU_misc_RNA_59 | 1.211348333 | 0.192298838 | | | 0 | | | |
| 22 | BSU_28260 | leuC | 1.20879374 | 0.000917827 | biosynthesis of leucin | 3-isopropylmalate de | 2 | Biosynthesis/ acquisit | Phosphorylation on an Ar | |
| 23 | BSU_22849 | ypzH | 1.157034029 | 0.301486735 | unknown | unknown | 1 | Newly identified sporulation proteins (based on | | |
| 24 | BSU_11290 | yjaV | 1.115583996 | 0.241513723 | [[category\|Sporulatic | [[protein\|SigE]]-depe | 1 | Sporulation proteins/ other(FC:0.41 68%) | | |
| 25 | BSU_18899 | yozX | 1.099514197 | 0.317003651 | unknown | unknown | 1 | Proteins of unknown function(FC:-0.35 60%) | | |
| 26 | BSU_26380 | yqaB | 1.098258018 | 0.214665205 | unknown | similar to phage-relat | 1 | Skin element(FC:0.45 62%) | | |

Figure 25: Overview of the table in sheet named “Mapped”. Category information is shown per gene.

#### 8. Overview of output sheets

##### 8.1 Operons

Make tables:

- Mapped: for all genes, the fold-changes of the genes in its respective operon.

##### 8.2 Regulons

Spread plots:

- RegSpreadPlot: plot of fold-changes of all included genes in the analyzed regulons.
- RegSpreadTab: table of fold-changes of all included genes in the analyzed regulons.
- RegSpreadPlotTopnFC: plot of fold-changes of all included genes in the top n regulons based on average FC.
- RegSpreakPlotTopnP: plot of fold-changes of all included genes in the top n regulons based on average p-value.
- RegSpreadTabTopnFC: table of fold-changes of all included genes in the top n regulons based on average FC.
- RegSpreadTabnP: table of fold-changes of all included genes in the top n regulons based on average p-value.

Bubble plots:

- RegRankPlot: bubbleplot of all selected regulons.
- RegRankPlotBest_v1: bubbleplot of most relevant regulon activity of all selected regulons.

- RegRankPlotBest_v2: bubbleplot of fraction of genes that follow most relevant regulon activity of all selected regulons.
- RegRankTable: Mapping_details (see “make tables”) but only for the genes that are included in the analysis.

Tables:

- Mapped: overview of regulon information for each gene. For each regulon, the average fold-change of genes following most relevant regulon activity and fraction of genes that follow this activity are shown.
- Mapping_details: tables showing 1) regulon details without regularity; 2) regulon details when regulator is repressed; 3) regulon details when regulator is activated; 4) best scoring regulon activity.

8.3 Categories

Spread plots:

- CatSpreadPlot: plot of fold-changes of all included genes in the analyzed categories.
- CatSpreadTab: table of fold-changes of all included genes in the analyzed categories.
- CatSpreadPlotTopnFC: plot of fold-changes of all included genes in the top n categories based on average FC.
- CatSpreakPlotTopnP: plot of fold-changes of all included genes in the top n categories based on average p-value.
- CatSpreadTabTopnFC: table of fold-changes of all included genes in the top n categories based on average FC.
- CatSpreadTab10nP: table of fold-changes of all included genes in the top n categories based on average p-value.

Bubble plots:

- CatRankPlot: bubbleplot of all selected categories.
- CatRankPlotBest_v1: bubbleplot of most relevant category activity of all selected categories.
- CatRankPlotBest_v2: bubbleplot of fraction of genes that follow most relevant category activity of all selected categories.
- CatRankTable: Mapping_details (see “make tables”) but only for the genes that are included in the analysis.

Tables:

- Mapped: overview of category information for each gene. For each category, the average fold-change of genes following most relevant category activity and fraction of genes that follow this activity are shown.
- Mapping_details: tables showing 1) Category details without regulation; 2) Category details when genes are upregulated; 3) Category details when genes are downregulated; 4) best scoring Category activity.

#### 9. Other ways to use GINtool data

9.1 Volcano plot display

A popular way to display transcriptome data is by means of a Volcano plot where the p-value (X-axis) of genes is plotted against the fold change (Y-axis). Using the average fold-change and p-values in the tables in the "Mapped" sheet, Volcano plots can be generated for both regulons as well as functional categories.

9.2 Comparing different experiments

The calculated average fold-change values for regulons in the "Mapped" sheet can be used as a convenient way to compare transcriptome data between different experiments by, e.g., using a scatter plot displaying the average fold-change value for a regulon from one experiment against the average fold-change found in the other experiment.

#### 10. Calculations

10.1 Median and Median absolute deviation (MAD)

As a robust measure, instead of the mean and standard deviation, the median and median absolute deviation (MAD) are used. The median is less prone to outliers and returns the ‘middle’ value of the observed values after sorting (from low to high). The MAD is a robust approximation of the spread around the median value and is calculated as the median of the residuals with respect to the median value determined earlier.

10.2 Average p-values

The p-values do not follow a normal distribution and they first have to be transformed using a Fisher-Z transformation into z-values. The average of these z-values is then back-transformed resulting in the average p-value.
